# The role of oxidative stress in seed priming to improve germination and vigour

**DOI:** 10.1101/2022.06.05.494903

**Authors:** Zane D. Marks, James M. Cowley, Rachel A. Burton, Tina Bianco-Miotto

## Abstract

Seed priming improves germination, but responses vary with procedure and plant used, potentially from differential responses in oxidative pathways. This study targeted oxidative pathways in seed priming, using hydrogen peroxide (H_2_O_2_), antioxidant-related micronutrients (Zn, Se and Mn), or a combination, to modulate germination and plant growth characteristics of hemp and barley. Hemp tolerated higher H_2_O_2_ (1 M vs 0.125 M) and micronutrients (6-fold greater) concentrations than barley, with the combination treatment significantly increasing hemp germination but decreasing barley germination. Seed priming did not improve hemp germination under salt stress (200 mM NaCl), but the micronutrient treatment improved germination of barley under salt stress (100 mM NaCl). Histological staining showed that micronutrient-primed root tips accumulated less O_2_^-^ in both non-stressed and salt-stressed conditions. We assessed seed priming benefits if grown in soil by measuring plant quality in three-week-old plants potted post-priming, but most quality factors were not significantly improved, except barley where shoot zinc content increased. In summary, seed priming with H_2_O_2_ and/or micronutrients increased germination in hemp, but only micronutrients increased barley germination. Future work will continue optimising the priming methodology and further investigate the role of oxidative stress in the observed responses.

## Introduction

Germination consists of the stages of physiological change in the initiation of the plant’s lifecycle. Germination begins when internal or external factors reach an ideal threshold for the seed. Different factors are important for germination, but in all plants, water is crucial, jumpstarting the initial stages of germination. Imbibition or the mass uptake in water rehydrates cells within the seed, restarting metabolic and respiratory function^1,2^. As nutrient stores are rapidly mobilised, the antioxidant system reactivates to compensate for the increase in respiratory activity. These processes continue until the final stage of germination which is the emergence of the radicle from within the seed. There are many ways to influence the germination process to increase nutrient content, enhance stress tolerance, or simply increase germination.

Seed priming is the enhancement of germination through the conditioning of seeds with an external factor^3,4^. Whilst the traditional method has been hydropriming (priming with water), several other methods have been used. For example, in basmati rice, hydropriming increased shoot length by 42% while priming with selenium (60 μM sodium selenite) improved shoot length by 125%^5^. The benefits of these priming techniques have been used to ensure uniform germination in crops, development of abiotic stress resistance, increased germination and potentially increased crop yield^4^. Whilst there are several methods of priming (reviewed in Farooq et al.)^4^, the foci of this study are redox priming (priming with oxidising agents) and nutri-priming (priming with micronutrients like vitamins and minerals) because both have interactions with reactive oxygen species.

Reactive oxygen species (ROS) are a group of chemicals formed from oxygen (O_2_) and characterised by their highly reactive nature and the potential damage they cause to biomolecules within the cell. ROS are usually by-products of metabolic and respiratory pathways including photosynthesis^6^. Heightened levels of ROS lead to a state known as oxidative stress, where antioxidant systems are unable to sufficiently detoxify the amount of ROS produced, which can cause massive damage and even cell death. Different ROS species exist but the primary species is superoxide (O_2_^-^); many of the other species are formed in the process of its reduction, with one of these secondary molecules being hydrogen peroxide (H_2_O_2_). H_2_O_2_ is also a ROS that is formed through the reduction of superoxide anions by the enzyme superoxide dismutase^6,7^ but can also form spontaneously through electron leakage in the mitochondria^6^. H_2_O_2_ is a moderately reactive molecule which in high quantities can cause cellular damage^6,8^; however, it is surprising that priming with H_2_O_2_ improves germination in several economically important species including pepper^9^, eggplant^10^ and pea^11^. H_2_O_2_ has secondary functions as a signalling molecule, and in moderate quantities is known to break dormancy in seeds^12,13^. H_2_O_2_ also cross-talks with plant phytohormones with evidence that it catalyses germination in moderate quantities^14^. Heightened levels of H_2_O_2_ can also accelerate the breakdown of nutrient reserves within the seed, making germination less energetically demanding^13^. H_2_O_2_ has an essential role in seed germination, but levels of H_2_O_2_ are closely regulated by the antioxidant system, therefore reaching an optimum level and balance of ROS is likely to be key to redox priming.

Antioxidant micronutrients are elemental cofactors which antioxidant enzymes need for catalysing the reduction of ROS. Antioxidant enzymes, such as superoxide dismutase and ascorbate peroxidase use these elemental cofactors as proton donors in the reduction of superoxide anions^15^. These cofactors include iron, zinc, manganese, selenium, and copper. These cofactors are also toxic in high amounts, inhibiting germination, plant growth, and causing damage to normal cellular functions^16–18^. However, priming with optimal concentrations of these cofactors (categorised as nutri-priming) can increase crop yields^19^, chlorophyll content^20^, and nutritional content in plants^21^. Additionally, priming canola, camelina, and bitter gourd seeds with selenium has been shown to increase abundance of antioxidants and improves ROS scavenging activities^22,23^.

Studies support that redox priming and nutri-priming affect metabolic and respiratory pathways through the regulation of oxidative stress. This pathway is crucial to germination, but it is unknown how micronutrients and H_2_O_2_ interact within the seed priming process. There is emerging evidence that micronutrients, when used in combination, can synergistically enhance germination and plant vigour^24^. Under the same principle, it is possible that there is a synergistic effect when priming with micronutrients and H_2_O_2_ in combination. In this study, we investigated the effect of priming seeds with H_2_O_2_, micronutrients and their combination on germination and plant vigour in industrial hemp (*Cannabis sativa*) and barley (*Hordeum vulgare*). These two species were selected for their contrasting agricultural history and physiological differences. Barley is a monocot which has been heavily cultivated and studied internationally for its agricultural benefit, becoming a backbone to the Australian grains industry for decades. Hemp is a eudicot species which has a newly emerged focus in the agricultural field. As such, there is little literature regarding the species including possible factors impacting germination and the effect that seed priming may have on the species.

## Results

### Dose-response optimisation of seed priming agents

Visible germination for both *Cannabis sativa* (hemp) and *Hordeum vulgare* (barley) generally occurs within the 18 h seed priming period. This is determined by protrusion of the radicle from the seed coat by 1 mm, visible without a microscope. To determine optimal concentration of micronutrients and H_2_O_2_, to be used individually and in a combination treatment, dose-response priming trials were run (summarised in Table 1). Optimal concentrations were selected based on the highest dose that did not cause detrimental effects to germinability (Supplementary Fig. 1-28).

**Table 1.**
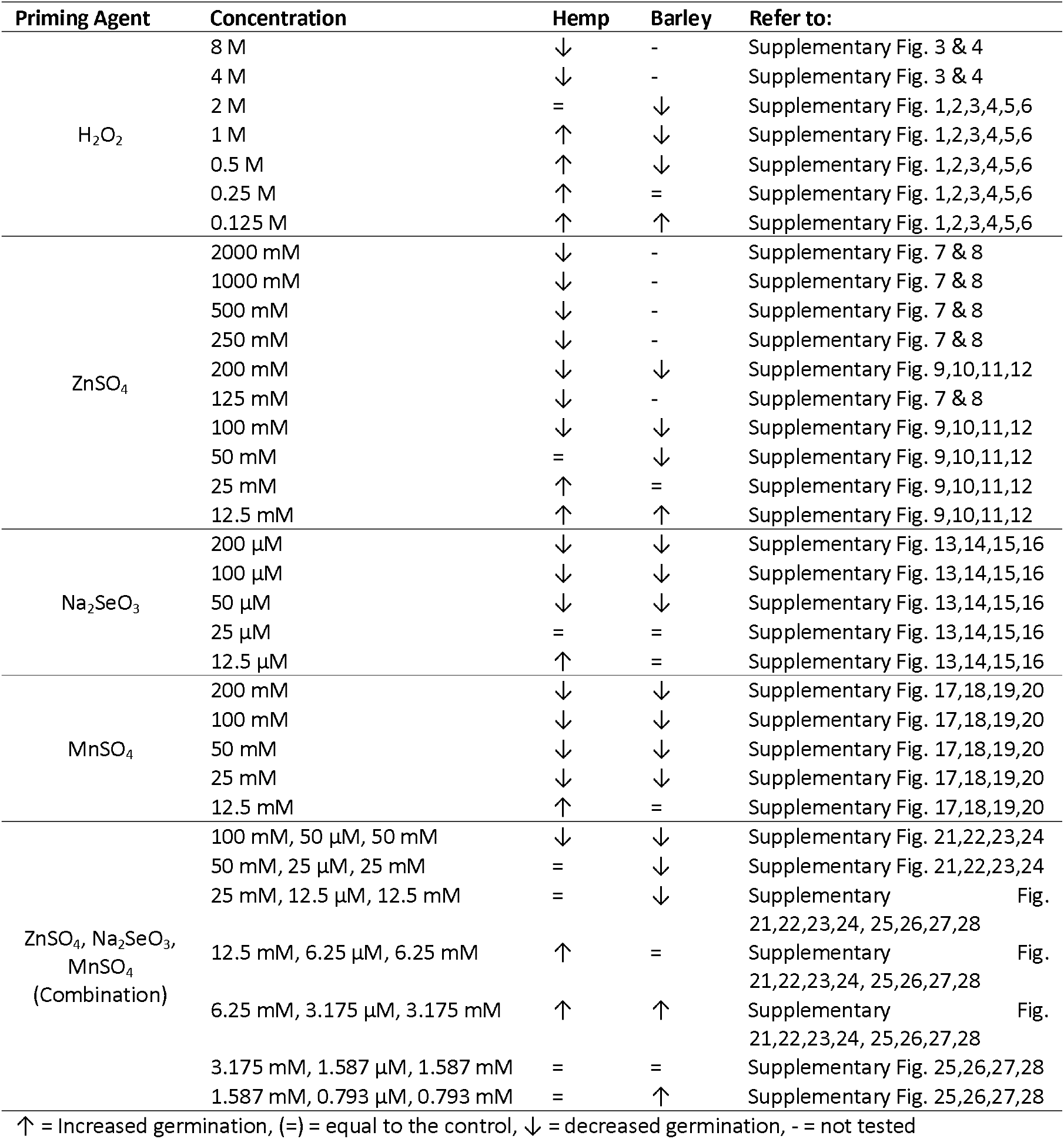
Summary of optimisation data addressing dose and effects seed for priming treatments used in the study

### Hydrogen peroxide

For hemp, hydrogen peroxide (H_2_O_2_) seed priming was effective in increasing rate of germination from the vehicle control (hydroprimed) in most concentrations (0.125 M, 0.25 M, 0.5 M, 1 M) (Supplementary Fig. 1-4). For barley, 0.125 M was the only treatment effective in increasing germination, with greater concentrations inhibiting germination (Supplementary Fig. 5-6). Interestingly, while 2 M H_2_O_2_ did not impact germination of hemp seeds, it entirely inhibited germination of barley (Table 1). For hemp, the optimal concentration selected was 0.5 M (Supplementary Fig. 1a) as this concentration was closer to the control at the initial time point of 0 h. At 24 h the germination percentage (relative to the control) was on par with less concentrated treatments, showing the greatest change in germination index (Supplementary Fig. 2). This effect in hemp is supported by relatively similar germination indices that the lower three dosages share (Supplementary Fig. 2). In barley, 0.125 M was selected as optimal due to noticeable increases in experiment 2 (Supplementary Fig. 5), and though experiment 1 shows a slight decrease at 48 h, this is not reflected in germination index where the value for the 0.125 M treatment was increased compared to the control (Supplementary Fig. 6).

### Zinc

Zinc, administered as zinc sulphate (ZnSO_4_), priming was effective at increasing the rate of germination in hemp at 25 mM and barley at 12.5 mM (Supplementary Fig. 7-12). All higher concentrations inhibited germination (Table 1). For hemp, 25 mM was selected as optimal despite 12.5 mM further increasing germination, as 25mM was comparable to the control. (Supplementary Fig. 7-8). The 12.5 mM dose was selected for barley as it was the only concentration that did not negatively affect germination (Supplementary Fig. 12).

### Selenium

Selenium, administered as sodium selenite (Na_2_SeO_3_), had a similar inhibitory effect on germination at higher concentrations for both hemp and barley, where only 12.5 μM increased germination in both species (Supplementary Fig. 13-16). In hemp, 12.5 μM was selected from the trials as it both increased germination percentage against the control and germination index (Supplementary Fig. 13, 14), all other concentrations reduced germination. In barley, 12.5 μM was also selected, it had a relatively similar effect to the control in both germination index and germination percentage, with no other concentrations observed to increase germination (Supplementary Fig. 15,16). An interesting note is that some barley seeds developed a pinkish hue towards the base of the radicle in higher doses (data not shown).

### Manganese

Manganese, administered as manganese sulphate (MnSO_4_), showed an inhibitory or comparable effect to the control regardless of concentration for both hemp and barley (Table 1). In hemp, both germination percentage and germination index of the vehicle control was not comparable to other trials which is indicative of interference (Supplementary Fig. 17, 18). The 12.5 mM dose was selected based off increased/comparable germination index, observable germination, and relevant literature^25^ as due to how germination index is calculated results may have been skewed. A similar response was observed with barley as the vehicle control had a varied germination rate across both experiments (Supplementary Fig. 19), so 12.5 mM was selected as it was closest to the control value in experiment 2 and increased germination percentage in experiment 1 (Supplementary Fig. 19, 20).

### Zinc, selenium, and manganese combination

After optimization with single micronutrients, the following concentrations were selected for both hemp and barley, 25 mM Zinc, 12.5 μM selenium, and 12.5 mM manganese as a baseline for combination optimization. Since the combination of the micronutrients may result in toxicity, combinations were optimized for each species (Supplementary Fig. 21-28). In barley, there was an observable decrease in germination except in lower concentrations of the micronutrient combination where it was comparable to the control (Supplementary Fig. 23, 24, 27, 28). Hemp was more tolerant to the micronutrient combination with greater increases in germination percentage and index before hitting a threshold where germination remained similar to the vehicle control (Table 1). After optimization, the combination selected for hemp was 6.25 mM ZnSO_4_, 3.125 μM Na_2_SeO_3_, and 3.125 mM MnSO_4_ on the basis that this combination had the highest germination index out of the trials performed (Supplementary Fig. 22). Similarly, in barley, the same combination concentration (6.25 mM ZnSO_4_, 3.125 μM Na_2_SeO_3_, and 3.125 mM MnSO_4_) was selected based off germination index where the indices were on par with the control or greater (Supplementary Fig. 24 and 28).

### The effect of optimized seed priming treatments on germination of hemp and barley

The combination priming treatment had an increased or comparable effect on germination than either agent alone (Fig.1). The H_2_O_2_ treatment had a comparable effect to the combination treatment whilst the micronutrient treatment increased germination but not to the same level as other treatments. This aligns with the optimisation trials for hemp, where H_2_O_2_ treatment demonstrated increased germination at most concentrations. For barley, the micronutrient treatment increased germination, while the combination and H_2_O_2_ inhibited germination relative to vehicle control (Fig. 2).

**Figure 1:**
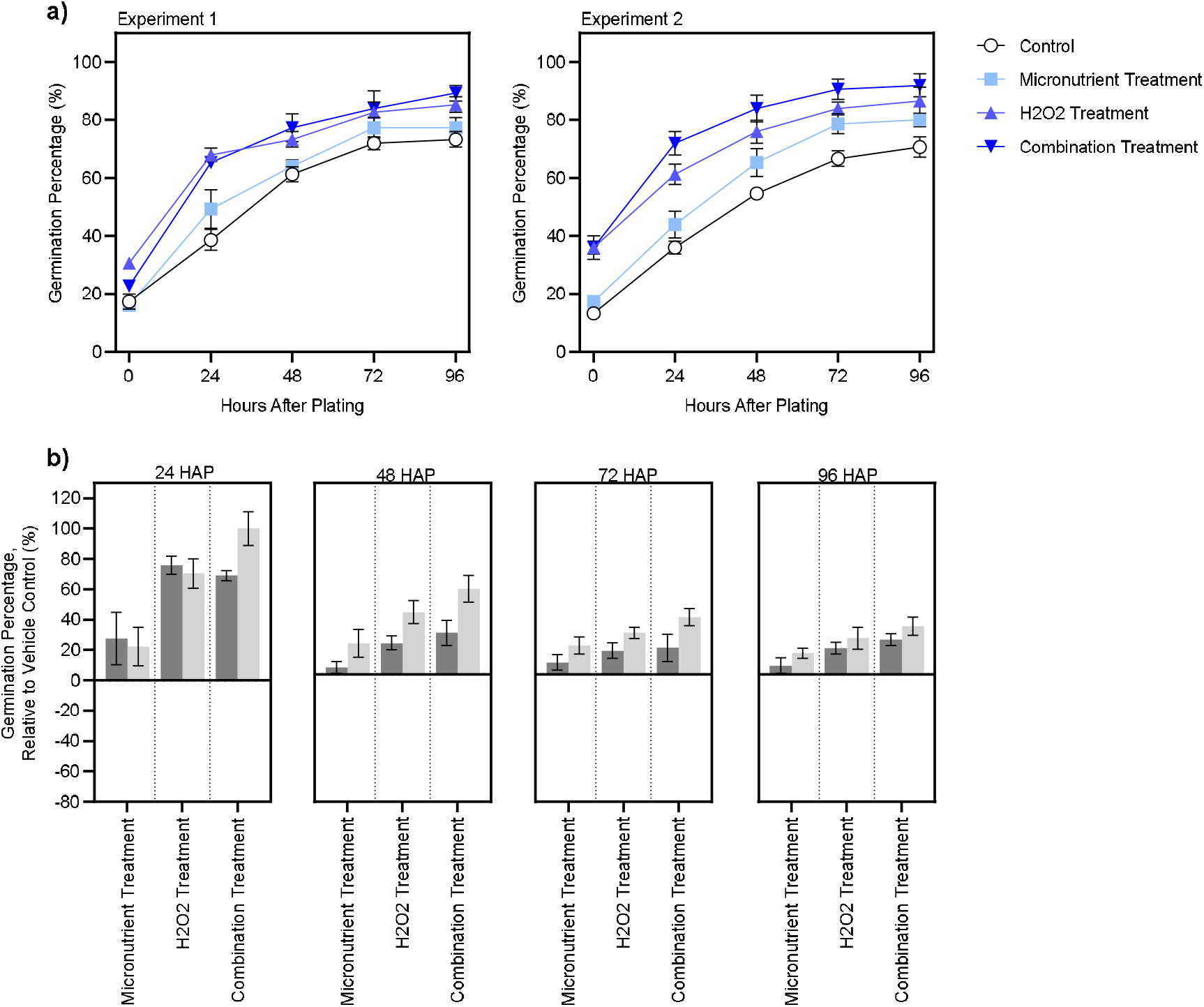
Hemp (*Cannabis sativa*) germination after optimised seed priming treatments a) Absolute germination percentage and b) germination percentage relative to the control (hydroprimed). Seeds were germinated in a 16/8-h light/dark controlled incubator after priming. Priming treatments included Micronutrient Treatment: 12.5 mM ZnSO_4_, 6.25 μM Na_2_Se_3_O, 6.25mM MnSO_4_; H_2_O_2_ Treatment: 500 mM H_2_O_2_ and Combination Treatment: 500 mM H_2_O_2_, 12.5 mM ZnSO_4_, 6.25 μM Na_2_Se_3_O, 6.25 mM MnSO_4_. The data shown is the mean of 3 replicates and standard error is shown as vertical bars. In (b), dark grey and light grey refer to independent experiment 1 and 2, respectively. In (b) HAP refers to “hours after plating”.

**Figure 2:**
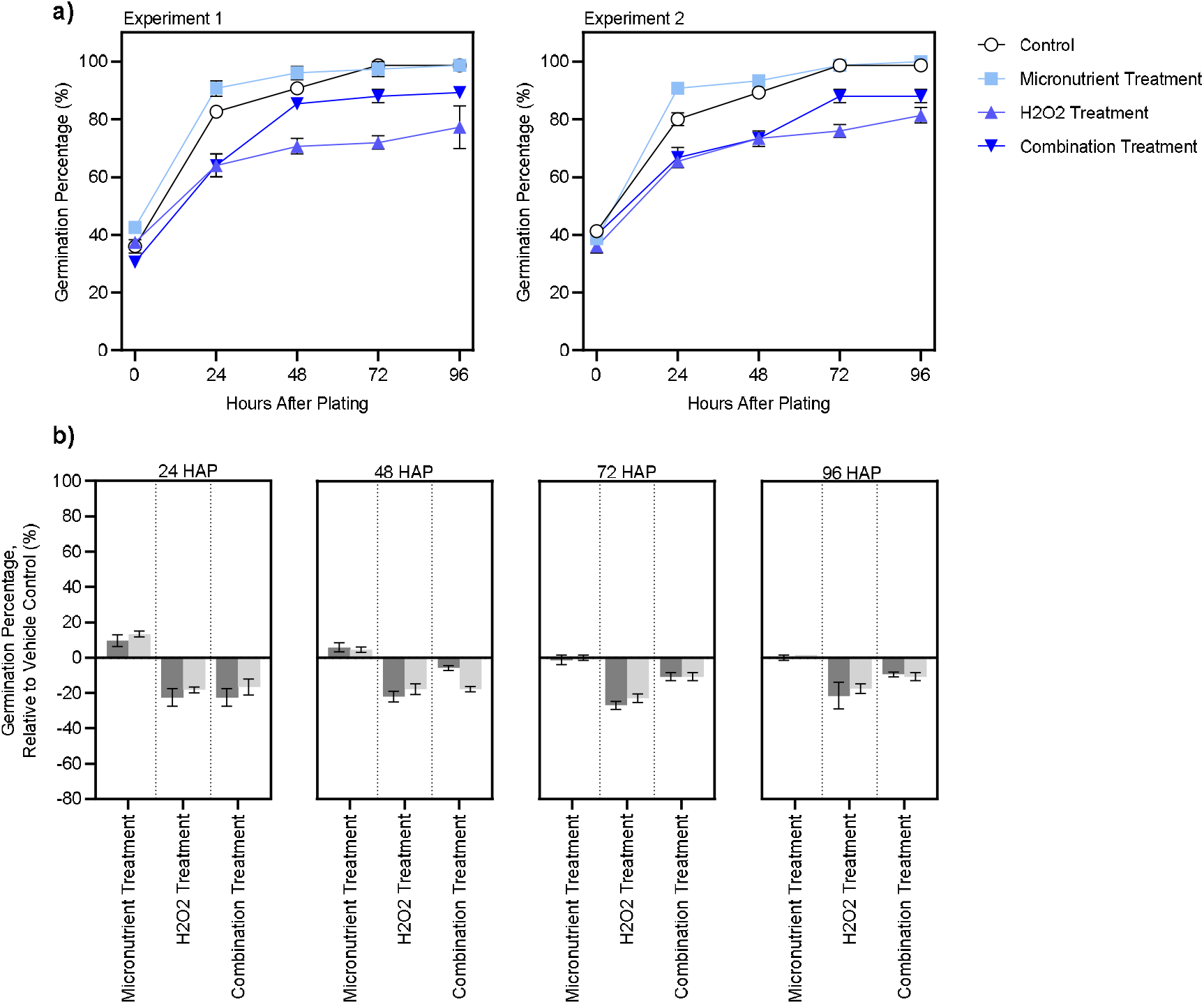
Germination of barley (*Hordeum vulgare*) after optimised seed priming treatments. a) Absolute germination percentage and b) germination percentage relative to the control (hydroprimed). Seeds were germinated in a 16/8-h light/dark controlled incubator after priming. Priming treatments included Micronutrient Treatment: 12.5 mM ZnSO_4_, 6.25 μM Na_2_Se_3_O, 6.25mM MnSO_4_; H_2_O_2_ Treatment: 500 mM H_2_O_2_ and Combination Treatment: 500 mM H_2_O_2_, 12.5 mM ZnSO_4_, 6.25 μM Na_2_Se_3_O, 6.25 mM MnSO_4_. The data shown is the mean of 3 replicates and standard error is shown as vertical bars. In (b), dark grey and light grey refer to independent experiment 1 and 2 respectively. In (b) HAP refers to “hours after plating”.

### Zinc and manganese levels in seeds primed with the micronutrients

Both hemp and barley displayed significantly (*p*<0.05) higher levels of both zinc and manganese when primed with the micronutrient treatment confirming that these micronutrients were taken up by primed seeds (Table 2). Under hydropriming (control) conditions it was found that hemp had a higher zinc and manganese level than barley. In hemp there was a 17.8-fold increase in zinc levels and a 3.8 fold increase in manganese levels when primed with the micronutrient treatment. Whilst barley demonstrated a 26.7-fold increase in zinc levels and a 15.2 fold increase in manganese levels under priming with micronutrients. Due to very low levels of selenium even in primed seeds, content was unable to be measured.

**Table 2.** ICP-OES analysis of hemp and barley seeds primed with the optimised micronutrient combination (12.5mM ZnSO_4_, 6.25 μM Na_2_Se_3_O, 6.25 mM MnSO_4_) compared to the control (hydroprimed). Values presented are means ± SE. FD = fold difference compared to control. **** = *p*<0.0001; *** = *p*<0.001; ** = *p*<0.01; comparing control and micronutrient treatments by Student’s *t*-test

|  | Hemp (*Cannabis sativa*) |  |  | Barley (*Hordeum vulgare*) |  |  |
| --- | --- | --- | --- | --- | --- | --- |
| Treatment | Control | Micronutrient | FD | Control | Micronutrient | FD |
| Zn (mg/kg) | 17.893<br>± 0.23 | 318.591<br>± 16.63**** | 17.8 | 4.993<br>± 0.05 | 133.37<br>± 15.55** | 26.7 |
| Mn (mg/kg) | 24.479<br>± 0.32 | 93.641<br>± 5.47*** | 3.8 | 2.309<br>± 0.04 | 35.025<br>± 4.18** | 15.2 |

### Germination of primed seeds under salt stress

In Fig. 3 the effects of salt stress (200 mM NaCl) on germination of hemp seeds is shown. There was an observable decrease in germination in salt stressed seeds when compared to non-stressed seeds (Fig. 3a). Both the micronutrient and combination treatment somewhat alleviated the effect of the stress stimuli, with the micronutrient treatment showing the greatest change (Fig. 3b). Conversely H_2_O_2_ had little effect on salt stressed germination. Fig. 3c highlights differences in germination index of salt stressed treatments compared to the control. Whilst there was an observable increase in all treatments relative to the control in experiment 1, there was no statistically significant difference (*p*>0.05) between treatments under stress conditions. The effects of priming and stress on the germination index of hemp seeds was tested using a two-way ANOVA. While priming and stress significantly affected germination index (*p<0*.*05)*, there was no significant interaction (*p>*0.05) indicating that seed priming did not alter the effects of the salt stress or *vice versa*. Further to this, no seed priming treatment significantly improved the germination index of salt stressed hemp seeds compared to the stressed control.

**Figure 3:**
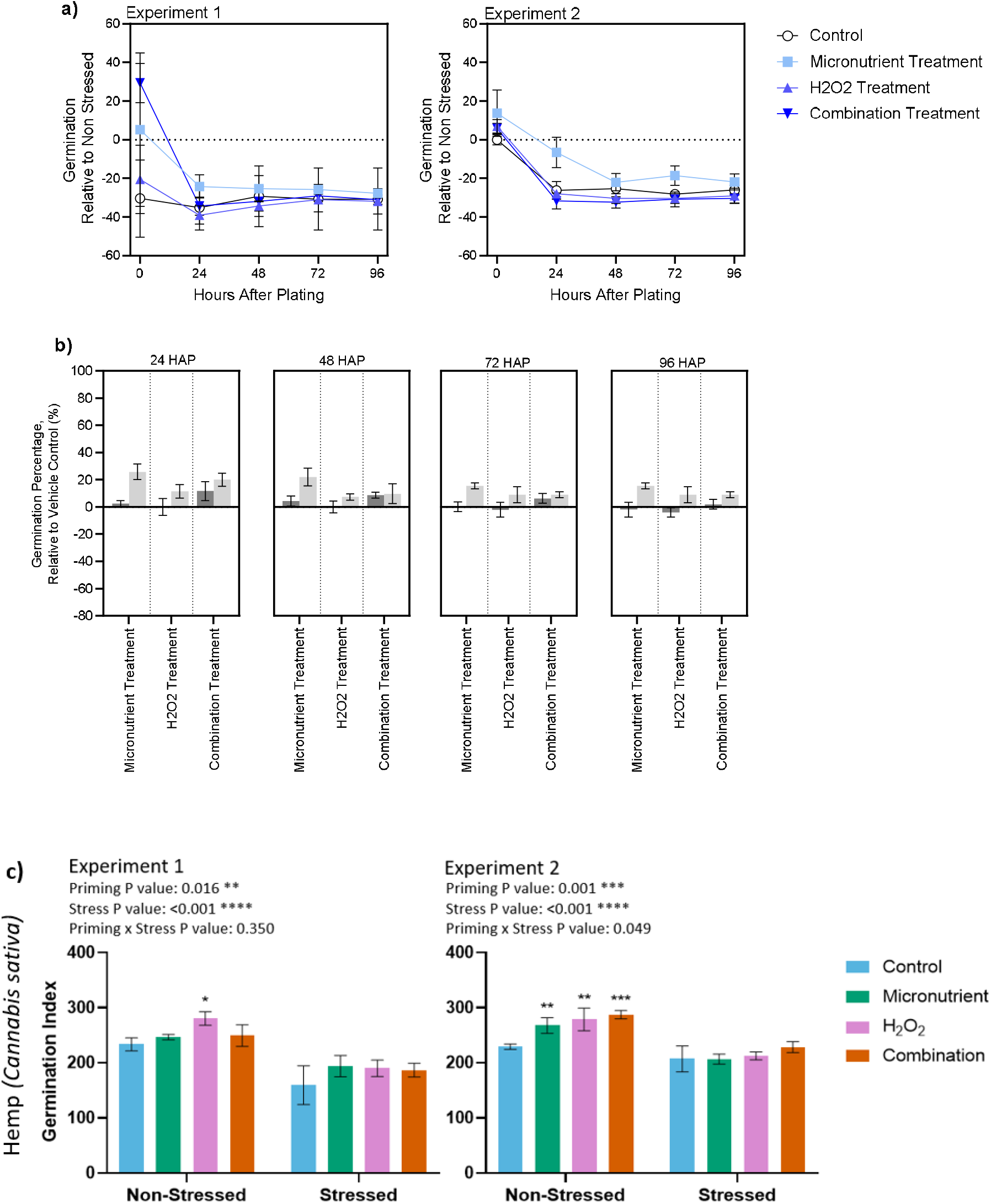
Effect of priming on response to salt stress in hemp (*Cannabis sativa*). a) Germination relative to non-stressed, b) germination percentage relative to the control (hydroprimed) and c) germination index of hemp seeds germinated in a 16/8-h light/dark controlled incubator primed with different treatments and germinated in 200 mM NaCl solution. Priming treatments included Micronutrient Treatment: 12.5 mM ZnSO_4_, 6.25 μM Na_2_Se_3_O, 6.2 5mM MnSO_4_; H_2_O_2_ Treatment: 500 mM H_2_O_2_ and Combination Treatment: 500 mM H_2_O_2_, 12.5 mM ZnSO_4_, 6.25 μM Na_2_Se_3_O, 6.25 mM MnSO_4_. The data shown is the mean of 3 replicates and standard error is shown as vertical bars. In (b), dark grey and light grey refer to independent experiment 1 and 2 respectively. In (B) HAP refers to “hours after plating”. *** = *p*<0.001; ** = *p*<0.01; * = *p*<0.05.

Fig. 4 shows the effect of salt stress (100 mM NaCl) on germination and germination index of barley seeds. There was a small decrease in germination in response to stress with the greatest decrease in germination seen at 24 h (Fig. 4 a, b). In barley, the germination index significantly decreased (*p*<0.05) in the combination and H_2_O_2_ priming treatment under both stressed and non-stressed conditions (Fig. 4c). Conversely the micronutrient priming treatment significantly increased (*p*<0.05) germination under stressed and non-stressed conditions (Fig. 4c). These trends were observable across both experiments. In barley, both the priming treatment and stress stimuli significantly impacted germination index (*p*<0.05). The interaction between the priming treatments and stress stimuli was statistically significant (*p*<0.05), indicative that seed priming impacted the effect of the stress stimuli.

**Figure 4:**
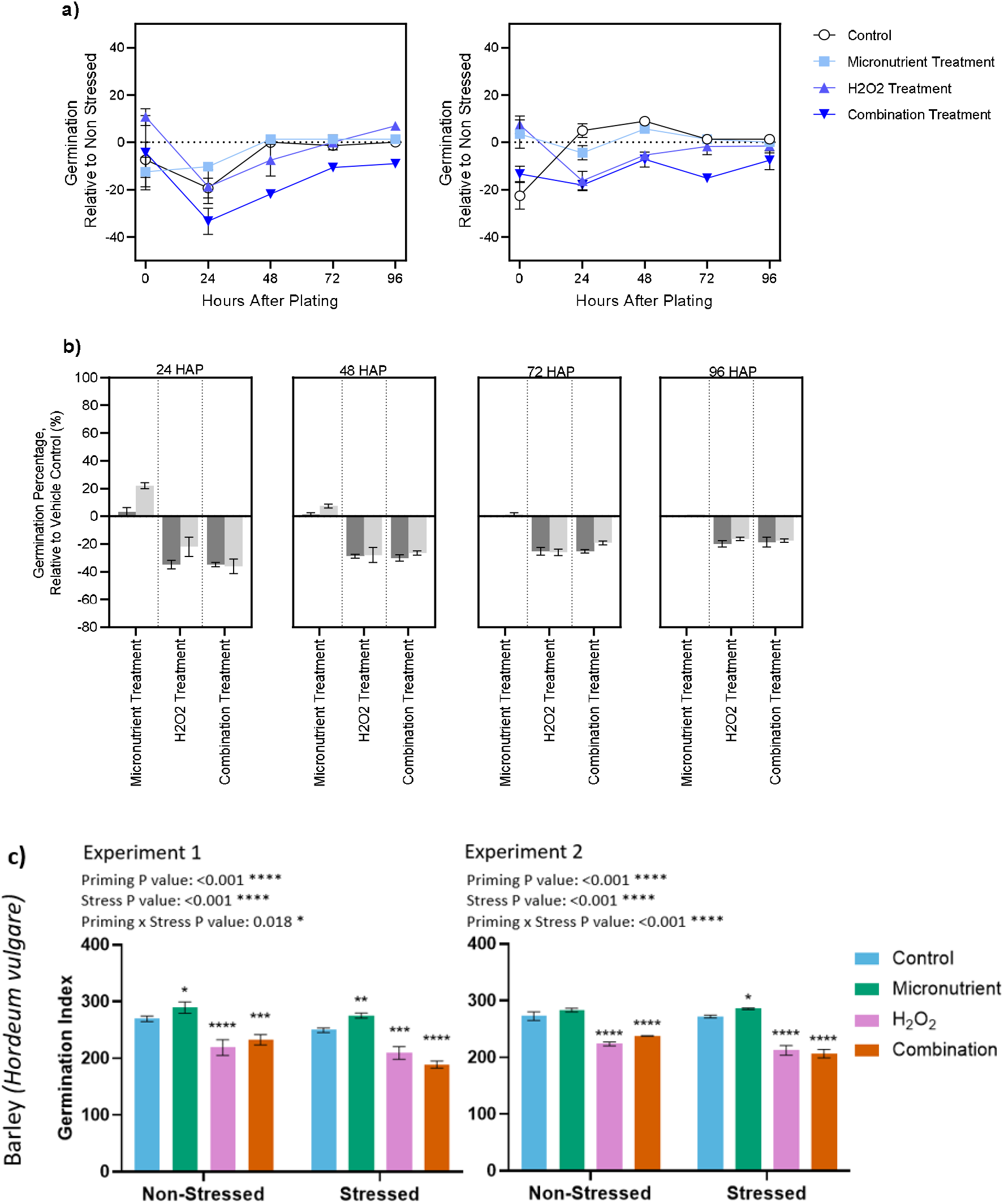
Effect of priming on response to salt stress in barley (*Hordeum vulgare*). a) Germination relative to non-stressed, b) germination percentage relative to the control (hydroprimed) and c) germination index of barley seeds germinated in a 16/8-h light/dark controlled incubator primed with different treatments and germinated in 100 mM NaCl solution. Micronutrient Treatment: 12.5 mM ZnSO_4_, 6.25 μM Na_2_Se_3_O, 6.25 mM MnSO4. H_2_O_2_ Treatment: 125mM H_2_O_2_. Combination Treatment: 125 mM H_2_O_2_, 12.5 mM ZnSO_4_, 6.25 μM Na_2_Se_3_O, 6.25 mM MnSO_4_. The data shown is the mean of 3 replicates and standard error is shown as vertical bars. In (B), dark grey and light grey refer to independent experiment 1 and 2 respectively. In (B) HAP refers to “hours after plating”. **** = *p*<0.0001; *** = *p*<0.001; ** = *p*<0.01; * = *p*<0.05

### Reactive oxygen species accumulation in radicles of primed seeds germinated with and without salt

Visualisation of superoxide (O_2_^-^) was conducted via NBT staining where increased presence of blue formazan chemical indicates a higher concentration O_2_^-^ anion and ROS activity as a proxy. Fig. 7 shows the difference in presence of O_2_^-^ with and without salt stress after seed priming in the two species. In non-stressed hemp, the H_2_O_2_ treatment led to the visually highest concentration of O_2_^-^ of all treatments, with the micronutrient treatment having the least. In stressed radicles of hemp seeds, a similar trend followed, with the micronutrient treatment (and the combination) visually having the least with nearly no NBT staining at the root tip and H_2_O_2_ having the most, along with some visible deterioration of the root surface. For barley, the non-stressed trial showed relatively similar coloration and concentration in all treatments except the H_2_O_2_ treatment. H_2_O_2_ visually had a higher concentration of blue formazan. In the barley stress trial, the micronutrient treatment also showed less ROS formation than its non-stressed equivalent and less than the control particularly in the root tip. Both the combination and H_2_O_2_ treatments did not produce enough material for the NBT assay under stressed conditions.

### Emergence and growth of primed seeds in soil

In the hemp potted growth experiment, the control (hydroprimed) had the highest successful emergence across all treatments and replicates with 77% (Fig. 6a). This was followed by the H_2_O_2_ and micronutrient combination treatment at 55%. Overall, it should be noted that over the three-week period that 1 plant from the H_2_O_2_ treatment (replicate 1) died, circumstances unknown, and that 2 plants died from the combination treatment (replicate 2), circumstances unknown. While these decreases in emergence were observable, none were determined to be statistically significant (*p*>0.05). In hemp, the micronutrient priming treatment led to the highest average dry weight by plant of 3.34 grams, while the lowest dry weight by plant was the combination treatment at 1.43 grams (Fig. 6b). The tallest average plant across the treatments was the combination treatment with 43.26 cm while the shortest on average was the H_2_O_2_ treatment with 38.62cm. The width of the plant is measured through the widest distance between two paired leaves. On average the widest plants were the combination treatment at 32.83 cm wide (Table 6b). Images representative of each treatment are shown in Supplementary Fig. 29. There was no statistically significant difference in zinc and manganese levels of fully grown hemp plants (Fig 6b). Whilst these additive differences in vigour were observable, none were determined to be statistically significant.

**Figure 5:**
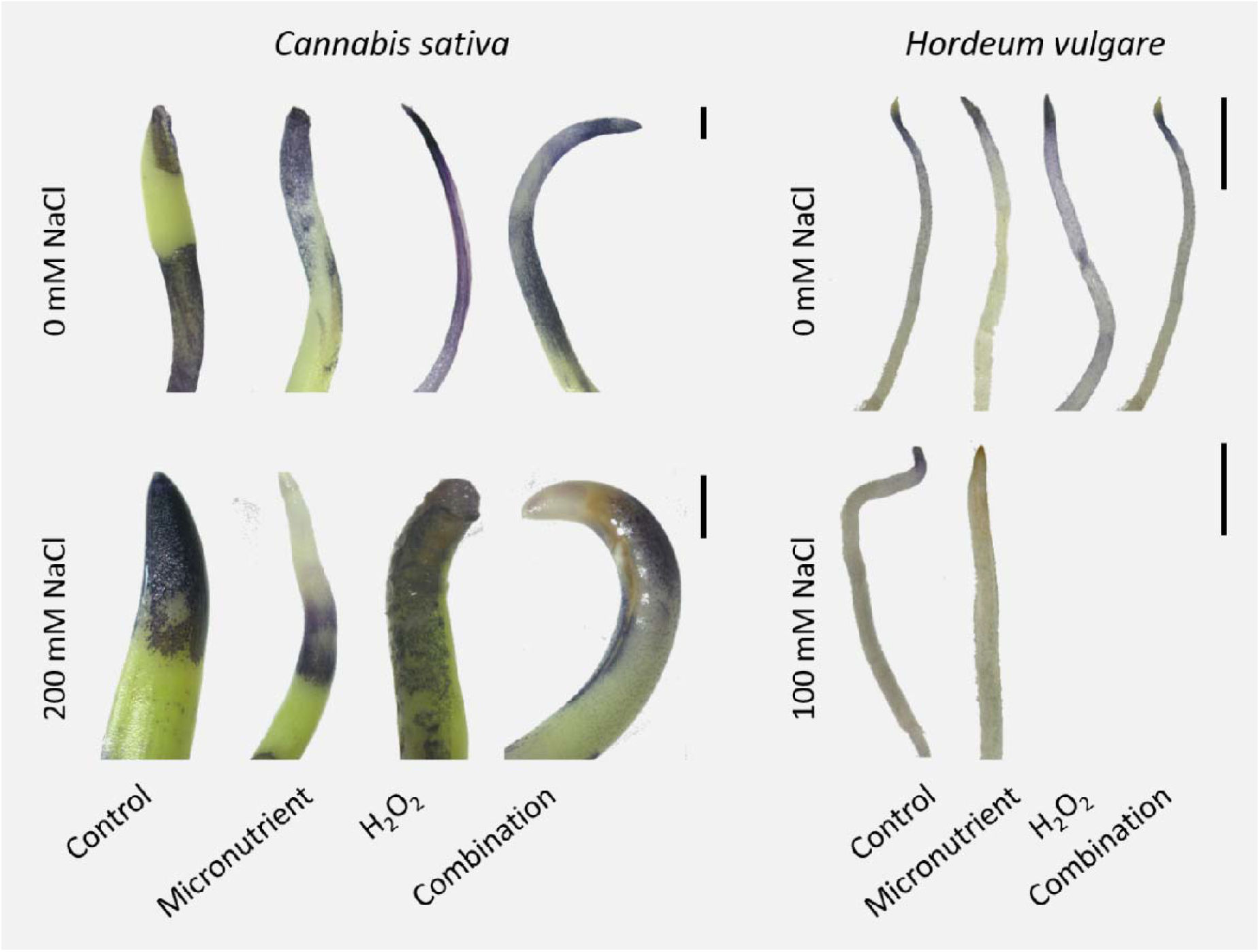
Visualisation of ROS activity in stressed and non-stressed hemp (*Cannabis sativa*) and barley (*Hordeum vulgare*). Localisation of superoxide (dark blue staining) using NBT staining of the radicle of 5-day old *Cannabis sativa* and *Hordeum vulgare* germinated without (0 mM NaCl) and with salt (200 mM or 100 mM NaCl for hemp and barley, respectively). Micronutrient Treatment: 12.5 mM ZnSO_4_, 6.25 μM Na_2_Se_3_O, 6.25 mM MnSO_4_. *C. sativa* H_2_O_2_ Treatment: 500 mM H_2_O_2_. *H. vulgare* H_2_O_2_ treatment: 125 mM *C. sativa* Combination Treatment: 500 mM H_2_O_2_, 12.5 mM ZnSO_4_, 6.25 μM Na_2_Se_3_O, 6.25 mM MnSO_4_. *H. vulgare* Combination Treatment: 125 mM H_2_O_2_, 12.5 mM ZnSO_4_, 6.25μM Na_2_Se_3_O, 6.25 mM MnSO_4_. Both the H_2_O_2_ and combination treatment did not produce enough material for NBT analysis therefore shown as blank. Scale for hemp = 100 μm; scale for barley = 1 cm.

**Figure 6:**
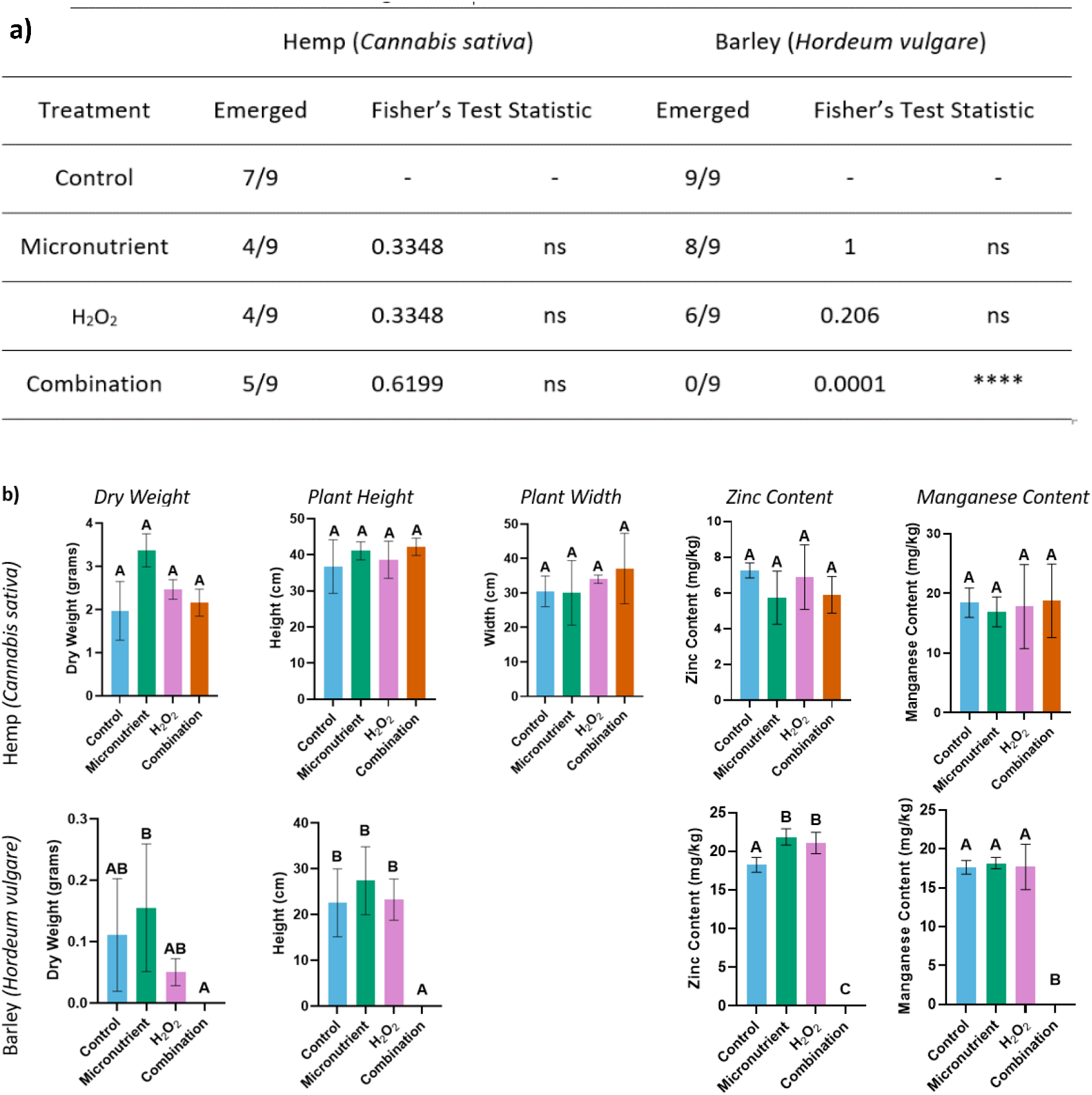
Impact of priming on potted growth, plant vigour and phenotypical factors in hemp (*Cannabis sativa*) and barley (*Hordeum vulgare*). a) Summary of Fisher’s Exact Test comparing emerged and non-emerged ratio between treatments and control. ns = not significant. b) The Dry weight, height, width, zinc and manganese content of three-week-old potted hemp and barley primed with different treatments. Hemp was grown in 16/8 hr light/dark photoperiod, 18/26 °C night/day-controlled environment room. Barley was grown in 16/8 hr light/dark photoperiod, 18/23 °C night/day-controlled environment room. Micronutrient Treatment: 12.5 mM ZnSO_4_, 6.25 μM Na_2_Se_3_O, 6.25 mM MnSO_4_. H_2_O_2_ Treatment: 500 mM H_2_O_2_. Combination Treatment: 500 mM H_2_O_2_, 12.5 mM ZnSO_4_, 6.25 μM Na_2_Se_3_O, 6.25 mM MnSO_4_. The data shown is the mean of 3 replicates and standard error is shown as vertical bars. Plant width is not shown for barley as it is not a relevant indicator for plant vigour in barley. **** = *p*<0.0001, ns = not significant, *p*>0.05.

In the barley potted growth trials, the combination priming treatment resulted in no emergence across all reps, which is represented as no data. There was no statistically significant difference between emergence of the remaining treatments and the control (Fig. 6a). The micronutrient treatment produced the highest dry weight, as well as the tallest height (Fig. 6b), but these increases were not statistically significant. It was observed that the H_2_O_2_ treatment in barley produced highly deformed leaves with a corkscrew shape, and the micronutrient treatment was observed to have undergone the tillering stage earlier than the control (Supplementary Fig. 30). There was determined to be a statistically significant increase in levels of zinc within three-week-old barley plants, when primed with the H_2_O_2_ and micronutrient treatments individually.

### Discussion

This study investigated the interaction between H_2_O_2_ and micronutrients when used as seed priming agents, and their impact on germination and plant vigour. The impact of seed priming with micronutrients and H_2_O_2_ varied greatly between hemp and barley, which was dependent on the concentration of the treatment. This is supported in the literature where in several plant species, high levels of micronutrients have been linked to increased levels of oxidative stress^26–28^. Conversely, optimised amounts of these micronutrients have been shown to increase germination, antioxidant content, and plant vigour factors^19,29^. A similar effect has been observed in ROS especially H_2_O_2_. Moderate quantities of H_2_O_2_ increase germination and break seed dormancy through its role as a secondary signalling molecule^14^, whereas higher quantities of H_2_O_2_ cause cell damage and even cell death^8^.

Our results showed a significant increase in germination index across all hemp seed priming treatments, under non-stressed conditions. The combination priming treatment significantly improved germination index which is notable as there is currently no published data on this combination of priming. As previously stated, both the H_2_O_2_ and micronutrient priming treatments have been shown to increase germination and various factors in plant growth^10,19,26,30^ including in hemp^31^. The results here support the notion that priming hemp seeds with H_2_O_2_ and micronutrients may produce a synergistic effect. This may be due to the treatments altering the balance of H_2_O_2,_ acting as signalling molecules, and micronutrients, mediating ROS damage, to improve germination. Conversely the potency of the H_2_O_2_ priming treatment in hemp is shown to be relative to that of the combination (Fig. 3c). This could lead one to speculate that the effect of H_2_O_2_ was so impactful in hemp that it alone was responsible for the changes seen in the combination priming treatment. Alternatively, a higher concentration of micronutrients may be required to conclusively show an additive or synergistic effect of H_2_O_2_ and micronutrients.

Whilst hemp showed a significant increase in germination index when primed with the combination treatment. The significant difference shown in the combination treatment was similar to the H_2_O_2_ treatment thus demonstrating that the combination treatment was comparable or better than either treatment alone (Fig 3). As the effect can only be relative to the effects on germination of other treatments using a measurement of gene expression levels or reduction in activity of antioxidant enzymes could provide useful insight into differences between the combination treatments and individual treatments.

We found that barley when primed with micronutrients, under both stressed and non-stressed conditions, demonstrated a significantly higher germination index compared to that of the control. This is in line with previous studies that highlighted the capability of micronutrients to increase germination and abiotic stress tolerance^32,33^. Zinc is well supported in the literature as having a role in increased chlorophyll production, photosynthesis, and antioxidant activity^32,34^. The micronutrient priming treatment increased levels of zinc in barley (Table 2) which has been shown to reduce the level of oxidative stress caused by high salinity^35^. However, this change in germination index of barley could also be attributed to the possibility that the plants weren’t under sufficient stress. This notion is supported in Fig. 4 which shows minimal levels of germination reduction when compared to non-stressed plants (particularly when compared to the reduction in stressed hemp; Fig. 3). In barley, both H_2_O_2_ and combination priming treatments significantly reduced germination index, both under stressed and non-stressed conditions. This is notable as concentrations of H_2_O_2_ were selected on the criterion that germination was not negatively impacted. Previous studies have highlighted a beneficial effect of H_2_O_2_ at certain levels and a toxic effect when a certain capacity or threshold is reached^36^. It could be speculated that the level of oxidative stress caused by both the H_2_O_2_ and combination priming treatment was toxic to barley. A study by Velarde-Buendia et al. highlights that in barley, ROS activity increases sensitivity to salt stress. High levels of ROS activate mediated potassium pumps causing an efflux of potassium which results in a cytotoxic effect^37^. ROS production from salinity when compounded with H_2_O_2_ could have intensified oxidative stress, overwhelming the antioxidant system and leading to death in barley.

It was found that micronutrient priming treatments resulted in reduced levels of ROS in the radicles of hemp and barley. This result is consistent with past literature that discusses the increase of elemental cofactors to mitigate ROS generation and accumulation in plants^38^. These results are also supported by a study by Singh et al. where radicles of *Vigna radiata* showed reduced accumulation of ROS when primed with copper chloride^39^ as copper is also an antioxidant cofactor. In addition, it was noted that in both the H_2_O_2_ and combination priming treatment of barley, not enough material was produced under salt stress for analysis. This may have been because, as previously mentioned, salt stress sensitivity is influenced by ROS which has been shown in previous studies to cause cytotoxic effects in salt stressed barley^36^. This, in combination with added oxidative stress from the H_2_O_2_ present in both treatments, could have overwhelmed the antioxidant capability of the seedling and in turn lead to increased damage to biomolecules and subsequent death^7^.

This study found that in hemp, regardless of priming treatment, there was no statistically significant difference in plant factors (dry weight, plant height, plant width, zinc, and manganese content). This result is not supported by the literature^19,29,35^. We can speculate that this effect may be due to the method of optimisation of our treatment. Treatments were optimised in a controlled environment without soil present, therefore the concentrations used may be inappropriate to produce an effect on hemp when grown in soil. In barley, the combination priming treatment resulted in no plant emergence in the potted growth trial. This could be due to soil composition, where the nutrient, salinity and/or biotic stress levels from the medium were varied from those of the optimisation trial. This could have resulted in an excess of micronutrients or salt which produced a toxic effect^40,41^. This combined with H_2_O_2_ might have conveyed a higher level of oxidative stress than the plant could handle causing mortality. This effect has been shown in previous studies such Reis et al where heightened levels of zinc caused plant mortality^20^. Complementing this was the lack of material from the combination treatment for the NBT, suggesting those differences in medium could amplify the toxicity of the treatment rather than being protective. When primed with micronutrients or H_2_O_2_, three-week-old barley was shown to have a significant increase in zinc content. Similar priming treatments have been shown to increase antioxidant capacity in other plant species such as wheat^25^ and bitter gourd^23^. This could account for the significant difference in zinc content.

In our study, hemp was shown throughout the results to be more tolerant to higher levels of micronutrients and H_2_O_2_ than barley. This could be due to naturally occurring variation between the two very different species. This may be supported by the native micronutrient content (Table 2) which shows that after hydropriming conditions, hemp seeds had a higher manganese and zinc content to barley. Due to the limited amount of literature on hemp there is only speculation into what physiological/genotypical differences could be the cause. Hypothetically, this could be due to difference in domestication status. A study by Han et al^42^ investigated differences in phenolic compound content of domesticated and wild barley, with wild accessions showing higher levels. As phenolics compounds are involved in stress and disease tolerance in plants^43^, the higher levels in wild accessions may be an adaptive difference to tolerate chronic oxidative stresses not experienced as frequently by cultivated barley. As hemp is not agriculturally domesticated it may retain many stress tolerance mechanisms associated with a wild phenotype.

It should be acknowledged that the results of this study encountered certain limitations. Whilst the study covered a variety of phenotypical measurements of hemp and barley, gene expression responses were not investigated. H_2_O_2_ has a known role as a regulatory signalling molecule for the expression of certain genes during germination^11,13^. Measurement of gene expression could clarify interactions between H_2_O_2_ and the expression of antioxidant enzymes that use elemental cofactors. As has been shown in this study there was great variation in the tolerance of H_2_O_2_ and micronutrients between hemp and barley. Literature also suggests an even greater variation in stress tolerance between plant species therefore the findings here may not be generalisable to other plant species^9,22,28,44^. This suggest that for different species of plants further optimisation would be necessary to convey the benefits of priming and limit the potential toxicity. As a note, during priming, many hemp and barley seeds germinated, exposing the radicle to the priming solution. This may have impacted growth factors as the radicle is a major site of respiratory activity, and exposure to the priming solutions can limit oxygen availability^45^.

## Conclusions

In this study we investigated the effect of H_2_O_2_ and micronutrient priming on germination and plant vigour. In hemp, all priming treatments significantly improved germination. Alternatively, in barley the micronutrient treatment increased germination while other priming treatments had an inhibitory effect. Both hemp and barley displayed reduced levels of ROS activity in the radical after being primed with micronutrients. Our results suggest that the combination priming treatment had a comparable or increased effect than individual treatments in hemp. Whereas in barley further optimisation would be required to fully establish the effect of the combination treatment. There was no statistically significant difference in plant vigour factors for hemp. Whist in barley the only statistically significant difference in plant quality was zinc levels. Future studies could investigate the efficacy of sequential (rather than simultaneous) priming with micronutrients and H_2_O_2_. In hemp, H_2_O_2_ was shown to increase germination rate. Therefore, an initial treatment of micronutrients followed by a drying stage, then a treatment of H_2_O_2_ could retain high micronutrient levels in fully grown plants and germination rate in seedlings. Additionally, a future direction could be to further optimise the priming treatments to account for the different composition of soil. In addition, further trials and optimisation could be run altering priming duration for each species to determine differential responses. Further investigation could be done into inheritability of the effects of various priming techniques. If beneficial the phenotypes are inherited by progeny, a series of growth and priming trials could be done to achieve simultaneous benefits from various priming methods.

In this study we have deepened our understanding into how micronutrients and H_2_O_2_ interact in oxidative pathways that impacts germination and plant vigour. This allows us to understand how beneficial phenotypical factors are formed and how to develop further priming techniques to increase efficiency and quality of the crops that we produce.

## Materials and Methods

### Plant materials

Seeds of barley (*Hordeum vulgare*) cv. Planet, a spring malt variety, were harvested from field trials conducted in Tarlee, SA and were kindly donated by Associate Professor Matthew Tucker (University of Adelaide). Seeds of hemp (*Cannabis sativa*) cv. Felina 32, a seed and fibre variety (<0.2% THC), were harvested from a field trial in Penola, SA and kindly donated by Good Country Hemp. Barley (*Hordeum vulgare*) seeds were stored dry at room temperature and hemp seeds were stored in sealed tubes at +4 °C.

The chemicals hydrogen peroxide solution 30%, (H_2_O_2_) (H3410, Sigma, Germany), zinc sulphate heptahydrate, ZnSO_4_.7H_2_O (Z0251, Sigma, Germany), manganese (II) sulfate monohydrate, MnSO_4_.H_2_O (M7899, Sigma, Germany), sodium selenite, Na_2_SeO_3_ (S5361, Sigma, Germany), 50 mM phosphate buffer, pH 6.8 (S0876, Sigma, Germany) and nitro tetrazolium blue chloride (NBT), C_40_H_30_N_10_O_6_·2Cl (N6876, Sigma, Germany) were acquired from Sigma Aldrich. The concentrations were diluted from stock concentrations in Milli-Q water and autoclave-sterilised.

### Priming procedure and germination assay

Using a pill counting tray, 25 seeds per replicate (3 replicates) were manually counted and placed into a 10 mL polypropylene centrifuge tube (PN 2024-07, Sarstedt, Germany). To ensure sterility, counted seeds in tubes were moved to a laminar flow hood with all tubes and instruments being sprayed with 70% Ethanol (Chemsupply, Australia). Seeds were disinfested by washing three times in sterilization solution (25% Undenatured Ethanol, 4% hypochlorite bleach (Cleera, Australia), 24% Milli-Q H_2_O, 1% Triton X-100 (C_14_H_22_O(C_2_H_4_O)_n_) (Sigma, Germany)). Tubes were inverted for 10 seconds, ensuring that the solution covered and washed all seeds. After the wash, the seeds were rinsed with autoclaved Milli-Q water three times before decanting as much liquid as possible from the seeds.

Five millilitres of autoclaved priming solution (prepared in Milli-Q water; vehicle control) to the sterilised seeds (approximately 1:5 w/v). Sealed tubes were removed from the laminar flow hood, placed into a test tube rack and the rack wrapped in aluminium foil to ensure that the samples were kept completely in the dark. The rack was placed on its side (tubes horizontal) on an orbital mixer to gently mix the seeds and priming agent for 18 h. After priming the tubes were moved back into the laminar flow hood for plating.

Primed seeds were removed from the priming solution and distributed on 20 mm x 100 mm petri dishes (664161, Grenier Bio-One, Austria) lined with 90 mm glass microfibre paper (PN1822-090, Whatman, United Kingdom) wetted with 5 mL of autoclaved water. Plates were sealed with parafilm and then transferred to an incubator under a controlled temperature (25 °C) in 16/8 light/dark cycle. At the relevant times after plating, germinating seeds were counted (radicle emergence >1 mm)

### Dose-response optimisation of priming agent concentrations

Optimisation experiments were performed for H_2_O_2_ and each micronutrient to determine the optimal concentration that improved germination (Table 1). H_2_O_2_ priming response was performed as serial dilutions with the range consisting of 2 M, 1 M, 0.5 M, 0.25 M, 0.125 M, and a vehicle control (0 M)^9,10^. A trial of higher concentration was conducted due to hemp’s positive response to H_2_O_2_, using the range of 8 M, 4 M, 2 M, 1 M (Supplementary Fig. 3,4).

For the zinc priming response, a serial dilution of 2 M, 1 M, 0.5 M, 0.25 M, and a control was prepared^18,19.^ The range was then reduced based on the initial trial, the range was adjusted to 200 mM, 100 mM, 50 mM, 25 mM, 12.5 mM.

In the selenium trial, the serial dilution range was adjusted in accordance to references in the literature^5,47,48^, the range consisted of 200 μM, 100 μM, 50 μM, 25 μM, 12.5 μM and the vehicle control.

Manganese priming response, the serial dilution range selected for this trial was based off the zinc trial, it consisted of 200 mM, 100 mM, 50 mM, 25 mM, 12.5 mM, and Control^24,49.^

Micronutrient combination were based off responses to the individual of the micronutrients. The first optimisation trial consisted of 5 treatments; Treatment 5: 100 mM ZnSO_4_, 50 μM Na_2_Se_3_O, 50 mM MnSO_4_, Treatment 4: 50 mM ZnSO_4_, 25 μM Na_2_Se_3_O, 25 mM MnSO_4_, Treatment 3: 25 mM ZnSO_4_, 12.5 μM Na_2_Se_3_O, 12.5 mM MnSO_4_, Treatment 2: 12.5 mM ZnSO_4_, 6.25 μM Na_2_Se_3_O, 6.25 mM MnSO_4_, Treatment 1: 6.25 mM ZnSO_4_, 3.125 μM Na_2_Se_3_O, 3.125 mM MnSO_4_. A second trial was conducted with different concentrations of the micronutrients, performed with 5 treatments; Treatment 5: 25 mM ZnSO_4_, 12.5 μM Na_2_Se_3_O, 12.5 mM MnSO_4_. Treatment 4: 12.5 mM ZnSO_4_, 6.25 μM Na_2_Se_3_O, 6.25 mM MnSO_4_. Treatment 3: 6.25 mM ZnSO_4_, 3.125 μM Na_2_Se_3_O, 3.125 mM MnSO_4_. Treatment 2: 3.125 mM ZnSO_4_, 1.5 μM Na_2_Se_3_O, 1.5 mM MnSO_4_. Treatment 1: 1.5 mM ZnSO_4_, 0.75 μM Na_2_Se_3_O, 0.75 mM MnSO_4._ From these trials, the optimised doses were selected: 12.5 mM ZnSO_4_, 6.25 μM Na_2_SeO_3_, 6.25 mM MnSO_4_ and different concentrations of H_2_O_2_ were used for barley and hemp, 125 mM, and 500 mM respectively.

### Micronutrient content analysis by inductively coupled plasma optical emission spectroscopy (ICP-OES)

Freeze-dried primed but ungerminated hemp and barley seeds and air-dried 3-week-old hemp and barley material were milled to a fine powder using a MM400 Mixer Mill (Restch, Germany). Ground powders were then digested and analysed by ICP-OES as per Wheal et al^50^.

### Germination in response to salt

The priming method was followed as described above for this trial, but the method was altered as the glass microfibre plates were instead wetted with a sterile NaCl solution (stressed) or water (vehicle control, non-stressed). Based on previous studies, the NaCl concentration administered for hemp was 200 mM and for barley was 100 mM NaCl^51,52^.

### Localisation of O_2_^.-^

Presence of superoxide (O_2_^-^) was detected using 0.5 mM NBT in 50 mM phosphate buffer pH 6.8 as modified from Singh et al^39^. Ten 5-day old samples were selected from each of the combination treatment, the micronutrient treatment, the H_2_O_2_ treatment and the controls, both under stressed and non-stressed conditions. The radicle was separated from the seed using a razor blade in the case of hemp and the roots in the case of barley. The roots were then placed in a 1.5 mL Eppendorf with the NBT solution for 25 min, after that the reaction was stopped via the removal of the NBT and rinsing the roots with distilled water. The stained material was then imaged under a stereo microscope fitted with a digital camera (Stemi 2000-C, Zeiss, Germany).

### Potted growth assay

To assess if the priming treatments affect vigour and emergence in soil conditions, a pot experiment was performed.

For barley, primed seeds (vehicle control, optimised H_2_O_2_, optimised micronutrients and combined H_2_O_2_ and micronutrients) were potted in coco peat. Three pot replicates were used with three seeds sown in a triangle formation at least 5 cm apart. Barley was grown in a controlled environment room with a 16/8 hr light/dark photoperiod and 18/23 °C night/day thermoperiod under high intensity discharge lamps.

For hemp, primed seeds (vehicle control, optimised H_2_O_2_, optimised micronutrients and combined H_2_O_2_ and micronutrients) were potted and grown in ‘goldilocks mix’ (50% UC Davis mix, 35% coco peat and 15% clay loam). Three pot replicates were used with three seeds sown in a triangle formation at least 5 cm apart and grown in a controlled environment room with a 16/8 hr light/dark photoperiod and 18/26 °C night/day thermoperiod under high intensity discharge lamps.

For both hemp and barley, emergence from soil was counted at 1-week post-potting. Each week for 3 weeks, a ruler was used to measure hemp and barley height and the width between two paired leaves for hemp. After 3 weeks of growth, plants were harvested from soil level and dried at 37 °C for five days before determining dry aboveground biomass.

### Statistical analysis

All germination experiments were performed with three plate replicates with two independent experiments.

Germination percentage relative to the vehicle control or Normalised percentage (NP), calculated as:

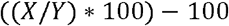

Where, at one time point, X is the treatment germination percentage and Y is the average of the corresponding control germination percentage.

Germination Index (GI), indicative of both velocity and percentage of germination, is calculated as per Kader M ^53^

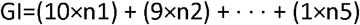

Where “n” is the number of seeds germinated on the respective day.

The germination index is calculated to represent the percentage and speed of germination in one directly comparable value, where the higher numeric value is indicative of increased speed and percentage. This method puts emphasis on the first days of germination rather than the later days

Statistical differences between groups (*p*<0.05) were determined by one-way Analysis of Variance (ANOVA) followed by a Duncan’s Multiple Range Test in GenStat 15^th^ Edition (VSI International, United Kingdom) or two-way ANOVA followed by a Bonferroni’s multiple comparison test in Prism 8.4.2 (Graphpad, United States).

A Fisher’s Exact Test (categorical) was performed to determine if there was a significant difference in the ratio of emerged and non-emerged seedlings between the vehicle control and primed seeds in the pot experiment(*p*<0.05).

Plotting was performed in Prism 8.4.2 (Graphpad, United States).

## Acknowledgements

The authors thanks Bogumila Tomczak and Michael McLaughlin from the University of Adelaide for performing ICP-OES analysis, Shi Fang (Sandy) Khor for technical assistance, Associate Prof. Matt Tucker for donation of barley seed and Good Country Hemp for the hemp seed donation.

