## Supplementary Figures for "The role of oxidative stress in seed priming to improve germination and vigour"

### *Supplementary Material:*

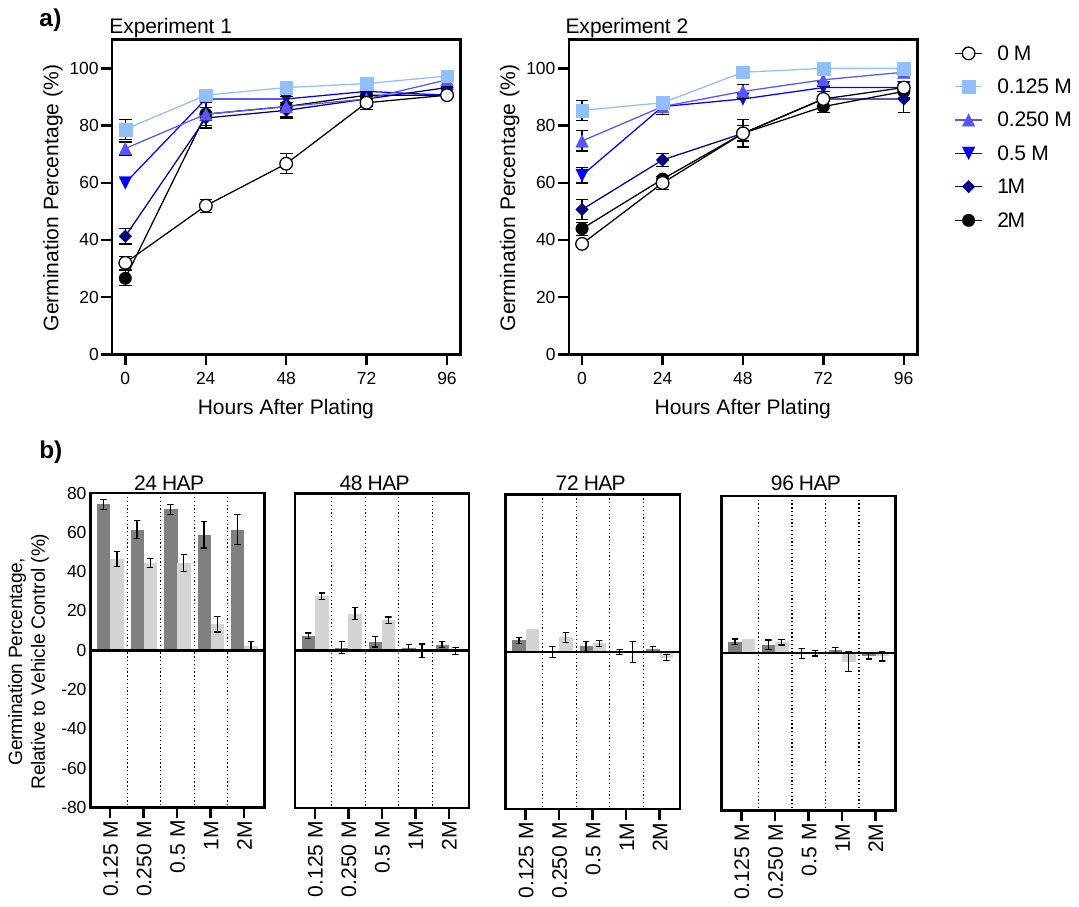

**Supplementary Figure 1:** Hydrogen peroxide (H_2_O_2_) priming optimisation experiments in hemp (*Cannabis sativa*).

a) Germination percentage and b) germination percentage relative to the control of hemp seeds germinated in a 25^o^C, 16/8 h light/dark controlled incubator for 96 h. The data shown is the mean of 3 replicates and standard error is shown as vertical bars. In (b), dark grey and light grey refer to independent experiment 1 and 2 respectively. In (b) HAP refers to “hours after plating”.

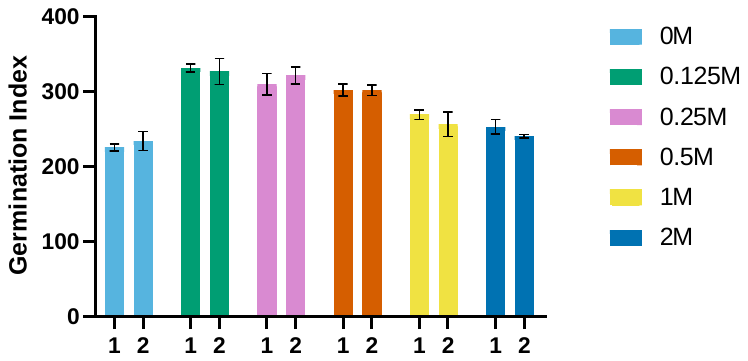

**Supplementary Figure 2**: Germination indices of hemp (*Cannabis sativa)* primed with varying concentrations of hydrogen peroxide (H_2_O_2_).

Hemp seeds were plated and grown in a 25^o^C, 16/8 h light/dark controlled incubator for 96 h. The data shown is the mean of 3 replicates and standard error is shown as vertical bars. “1” and “2” represent independent experiments done under the same conditions.

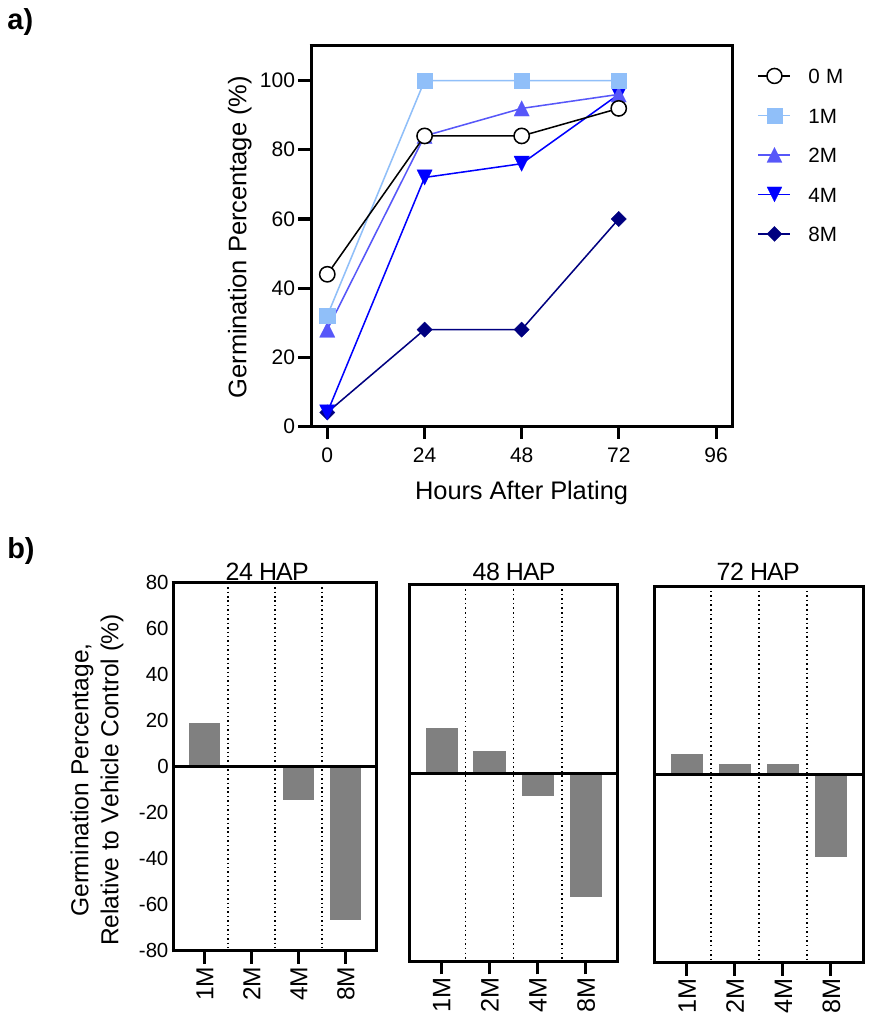

**Supplementary Figure 3**: Hydrogen peroxide (H_2_O_2_) priming optimisation experiments in hemp (*Cannabis sativa*).

a) Germination percentage and b) germination percentage relative to the control of hemp seeds germinated in a 25^o^C, 16/8 h light/dark controlled incubator for 72 h. The data shown is the result of one replicate. In (b) HAP refers to “hours after plating”.

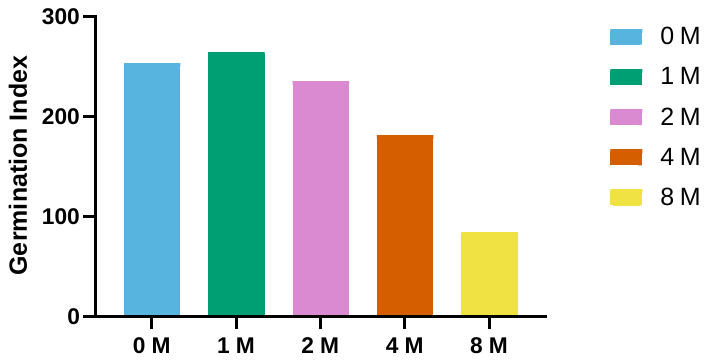

**Supplementary Figure 4**: Germination indices of hemp (*Cannabis sativa)* primed with varying concentrations of hydrogen peroxide (H_2_O_2_).

Hemp seeds were plated and grown in a 25^o^C, 16/8 h light/dark controlled incubator for 96 h. The data shown is the result of one replicate.

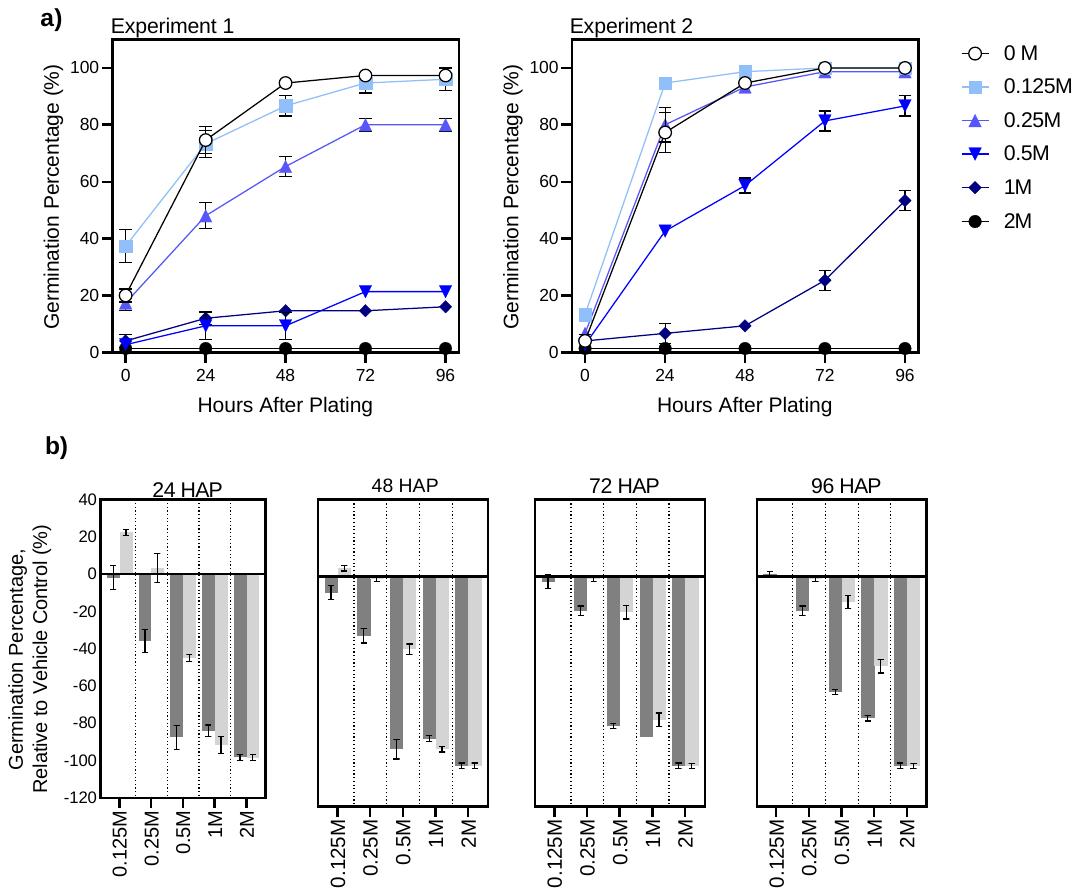

**Supplementary Figure 5**: Hydrogen peroxide (H_2_O_2_) priming optimisation experiments in barley (*Hordeum vulgare*).

a) Germination percentage and b) germination percentage relative to the control of barley seeds germinated in a 25^o^C, 16/8 h light/dark controlled incubator for 96 h. The data shown is the mean of 3 replicates and standard error is shown as vertical bars. In (b), dark grey and light grey refer to independent experiment 1 and 2 respectively. In (b) HAP refers to “hours after plating”.

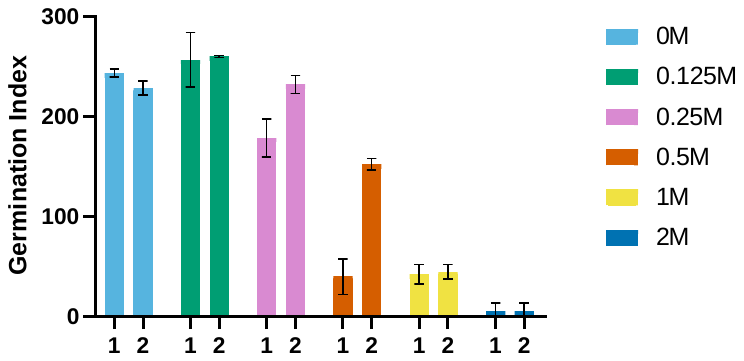

**Supplementary Figure 6**: Germination indices of barley (*Hordeum vulgare)* primed with varying concentrations of hydrogen peroxide (H_2_O_2_).

Barley seeds were plated and grown in a 25^o^C, 16/8-h light/dark controlled incubator for 96 h. The data shown is the mean of 3 replicates and standard error is shown as vertical bars. “1” and “2” represent independent experiments done under the same conditions.

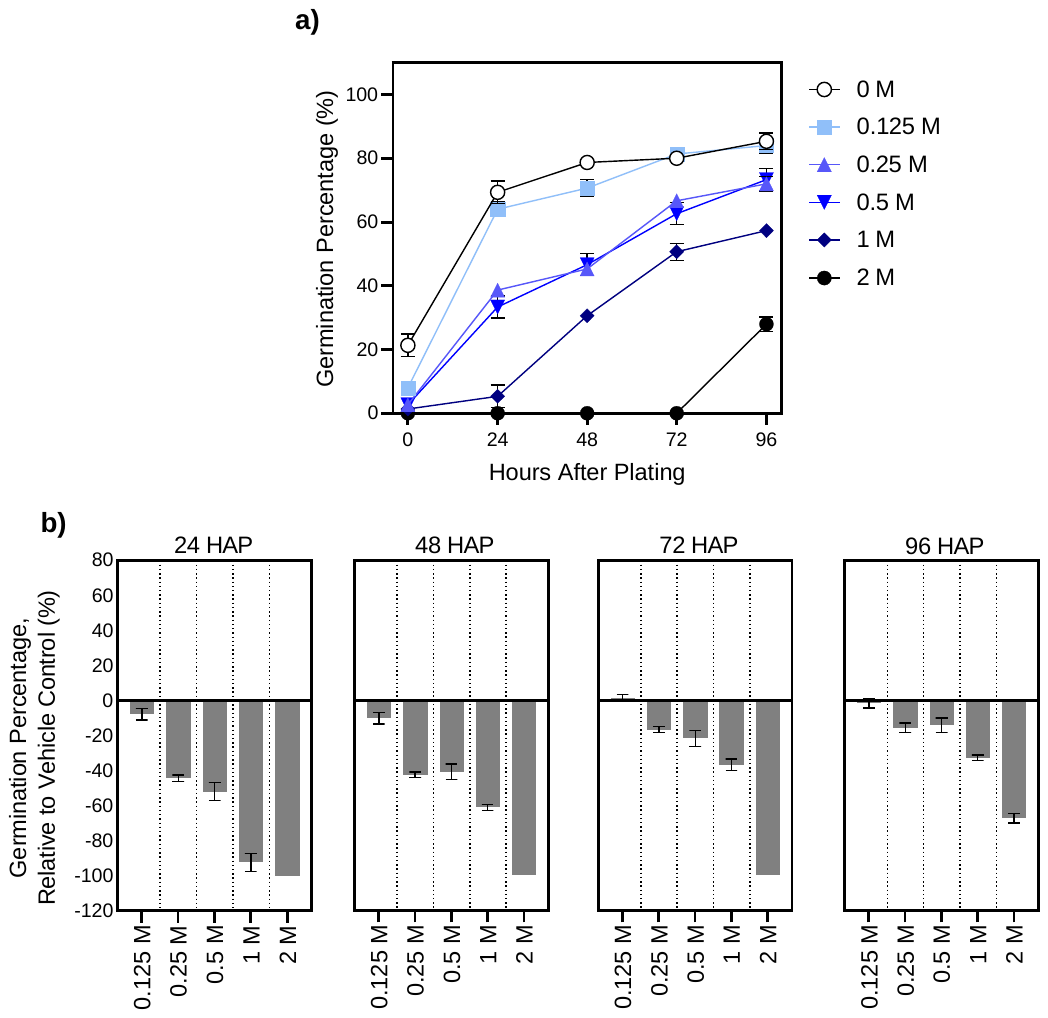

**Supplementary Figure 7**: Zinc (ZnSO_4_) priming optimisation experiments in hemp (*Cannabis sativa*).

a) Germination percentage and b) germination percentage relative to the control of hemp seeds germinated in a 25^o^C, 16/8 h light/dark controlled incubator for 96 h. The data shown is the mean of 3 replicates and standard error is shown as vertical bars. In (b) HAP refers to “hours after plating

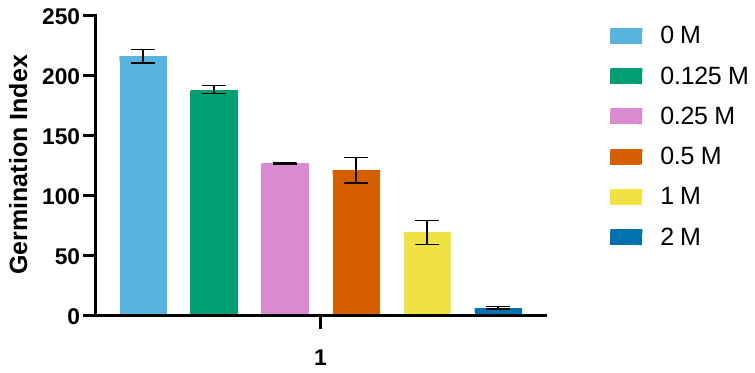

**Supplementary Figure 8**: Germination indices of hemp (*Cannabis sativa)* primed with varying concentrations of Zinc(ZnSO_4_).

Hemp seeds were plated and grown in a 25^o^C, 16/8 h light/dark controlled incubator for 96 h. The data shown is the mean of 3 replicates and standard error is shown as vertical bars.

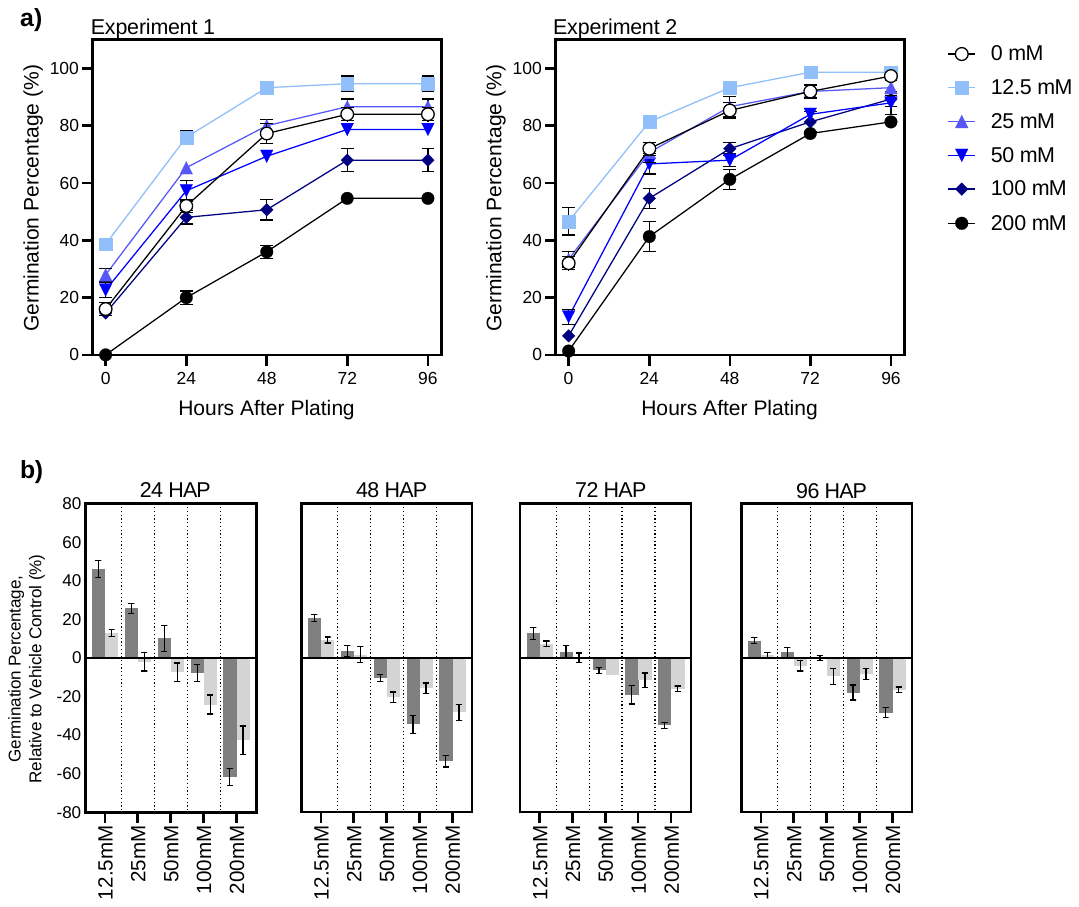

**Supplementary Figure 9**: Zinc (ZnSO_4_) priming optimisation experiments in hemp (*Cannabis sativa*).

a) Germination percentage and b) germination percentage relative to the control of hemp seeds germinated in a 25^o^C, 16/8 h light/dark controlled incubator for 96 h. The data shown is the mean of 3 replicates and standard error is shown as vertical bars. In (B), dark grey and light grey refer to independent experiment 1 and 2 respectively. In (B) HAP refers to “hours after plating”.

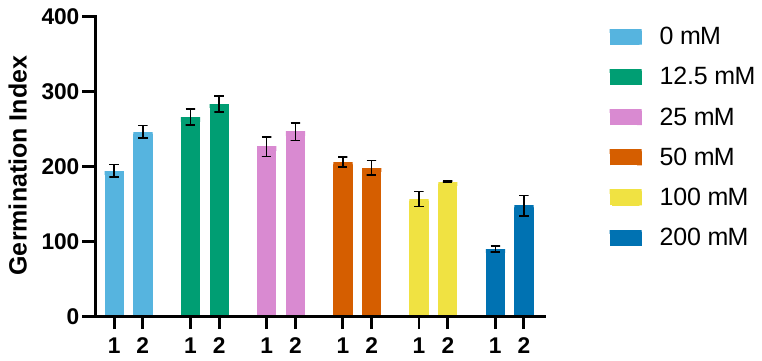

**Supplementary Figure 10**: Germination indices of hemp (*Cannabis sativa)* primed with varying concentrations of Zinc (ZnSO_4_).

Hemp seeds were plated and grown in a 25^o^C, 16/8-h light/dark controlled incubator for 96 h. The data shown is the mean of 3 replicates and standard error is shown as vertical bars. “1” and “2” represent independent experiments done under the same conditions.

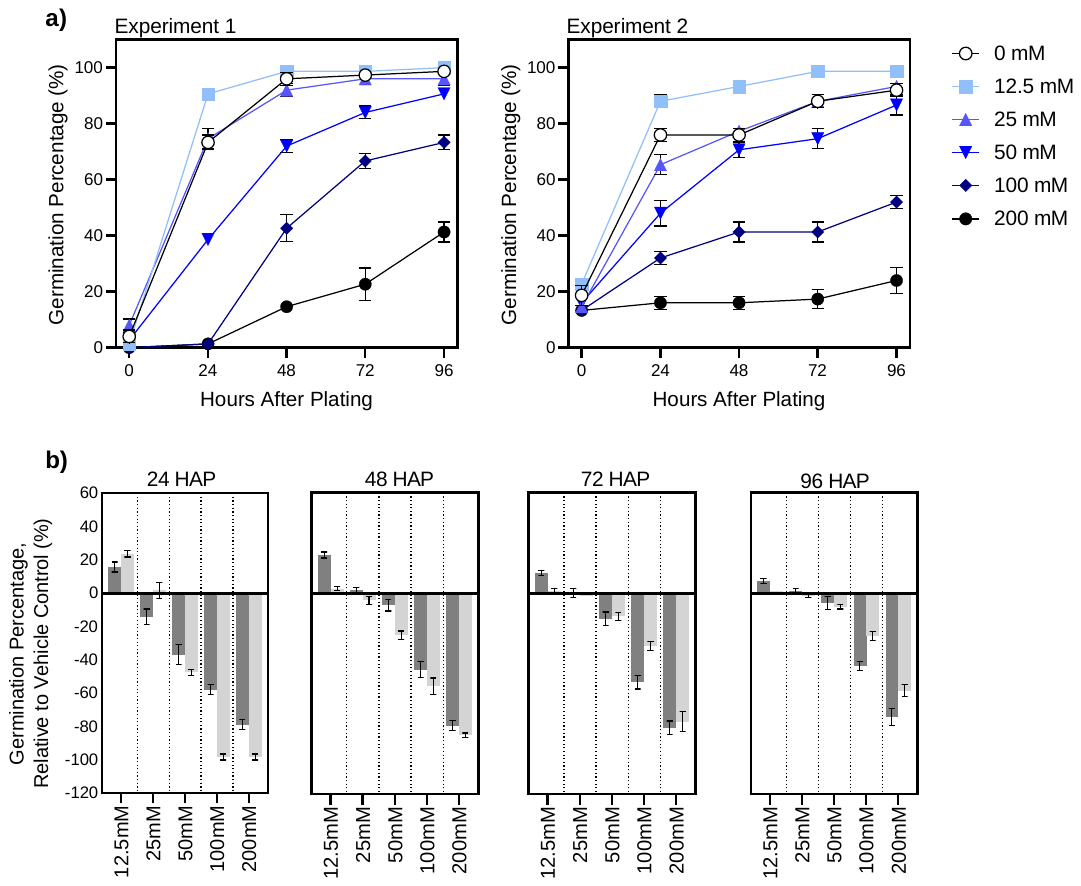

**Supplementary Figure 11**: Zinc (ZnSO_4_) priming optimisation experiments in barley (*Hordeum vulgare*).

a) Germination percentage and b) germination percentage relative to the control of barley seeds germinated in a 25^o^C, 16/8 h light/dark controlled incubator for 96 h. The data shown is the mean of 3 replicates and standard error is shown as vertical bars. In (b), dark grey and light grey refer to independent experiment 1 and 2 respectively. In (b) HAP refers to “hours after plating”.

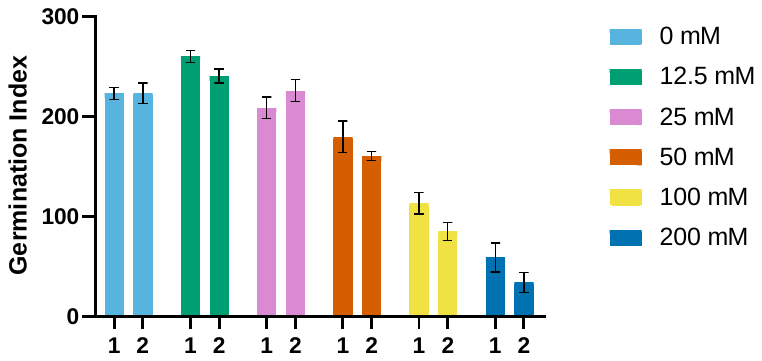

**Supplementary Figure 12**: Germination indices of barley (*Hordeum vulgare)* primed with varying concentrations of Zinc (ZnSO_4_).

Barley seeds were plated and grown in a 25^o^C, 16/8-h light/dark controlled incubator for 96 h. The data shown is the mean of 3 replicates and standard error is shown as vertical bars. “1” and “2” represent independent experiments done under the same conditions.

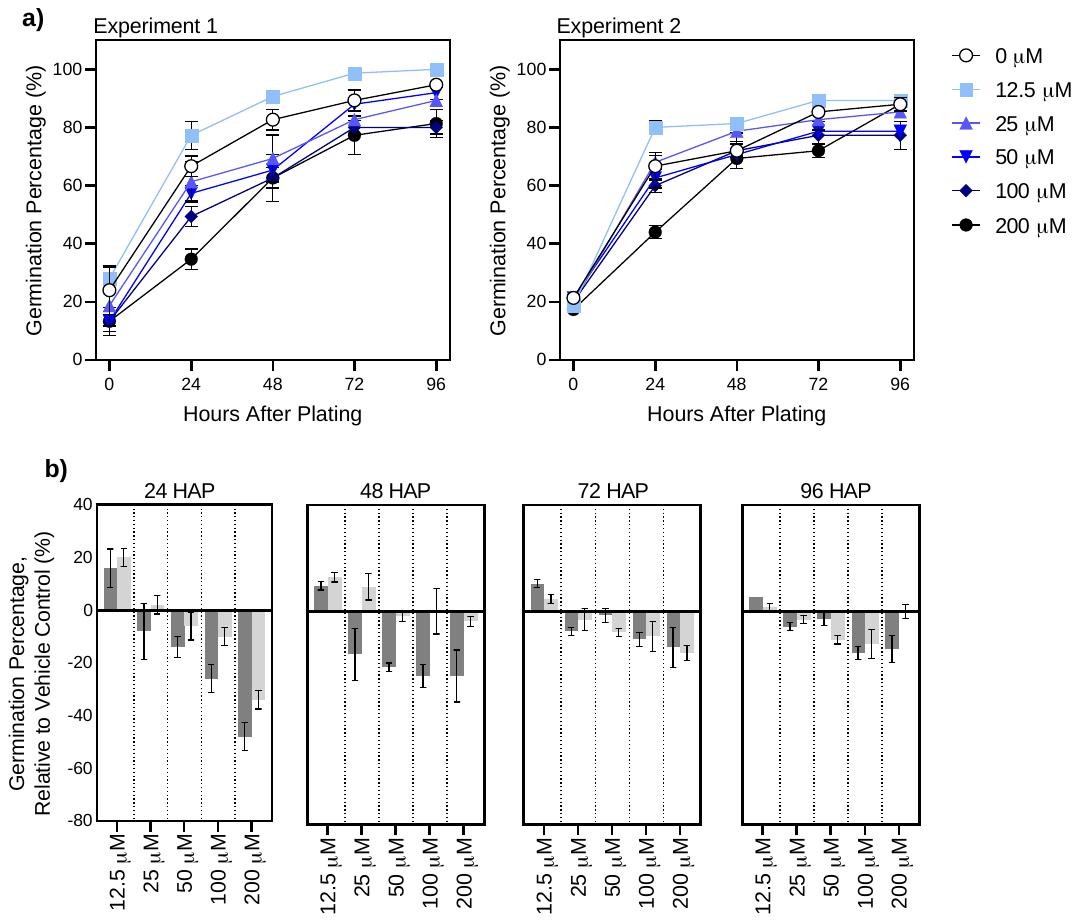

**Supplementary Figure 13**: Selenium (Na_2_SeO_3_) priming optimisation experiments in hemp (*Cannabis sativa*).

a) Germination percentage and b) germination percentage relative to the control of hemp seeds germinated in a 25^o^C, 16/8 h light/dark controlled incubator for 96 h. The data shown is the mean of 3 replicates and standard error is shown as vertical bars. In (b), dark grey and light grey refer to independent experiment 1 and 2 respectively. In (b) HAP refers to “hours after plating

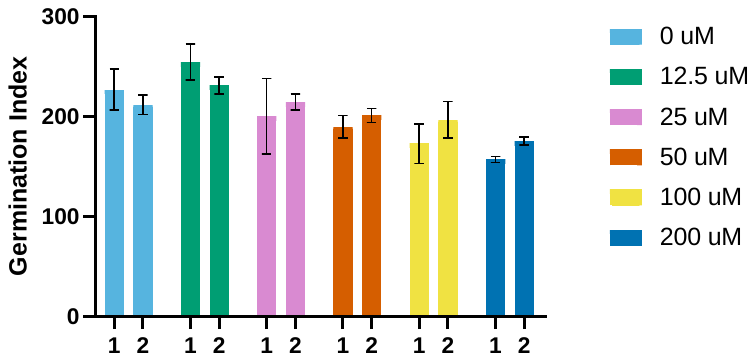

**Supplementary Figure 14**: Germination indices of hemp (*Cannabis sativa)* primed with varying concentrations of Selenium (Na_2_SeO_3_).

Hemp seeds were plated and grown in a 25^o^C, 16/8-h light/dark controlled incubator for 96 h. The data shown is the mean of 3 replicates and standard error is shown as vertical bars. “1” and “2” represent independent experiments done under the same conditions.

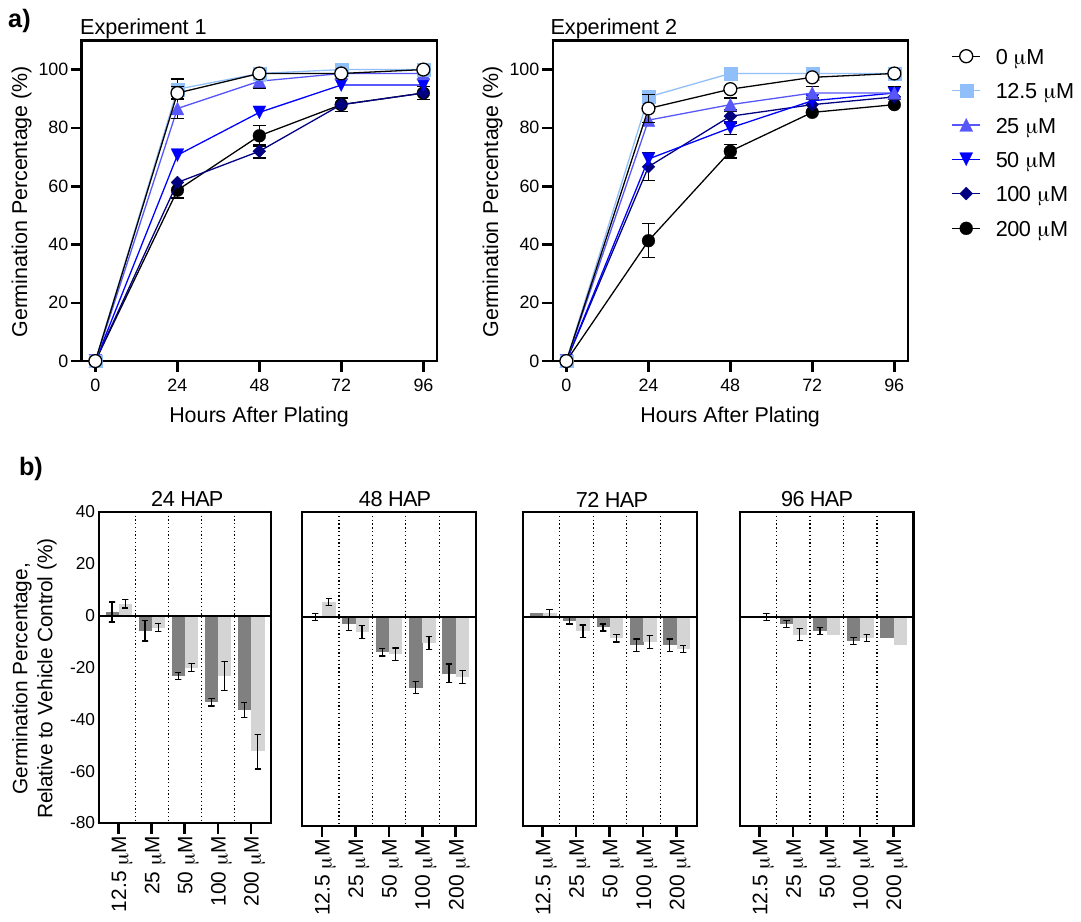

**Supplementary Figure 15**: Selenium (Na_2_SeO_3_) priming optimisation experiments in barley (*Hordeum vulgare*).

a) Germination percentage and b) germination percentage relative to the control of barley seeds germinated in a 25^o^C, 16/8-h light/dark controlled incubator for 96 h. The data shown is the mean of 3 replicates and standard error is shown as vertical bars. In (b), dark grey and light grey refer to independent experiment 1 and 2 respectively. In (b) HAP refers to “hours after plating”.

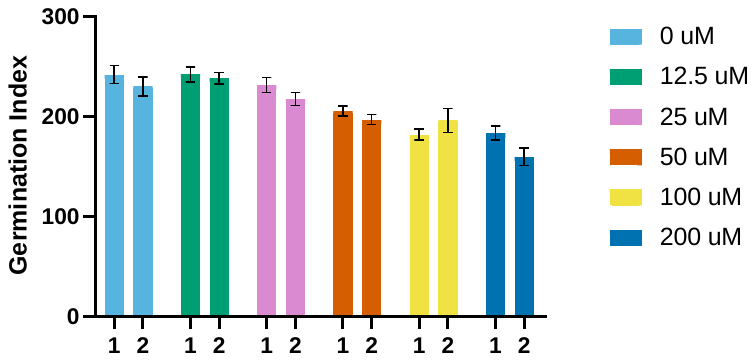

**Supplementary Figure 16**: Germination indices of barley (*Hordeum vulgare)* primed with varying concentrations of Selenium (Na_2_SeO_3_).

barley seeds were plated and grown in a 25^o^C, 16/8-h light/dark controlled incubator for 96 h. The data shown is the mean of 3 replicates and standard error is shown as vertical bars. “1” and “2” represent independent experiments done under the same conditions.

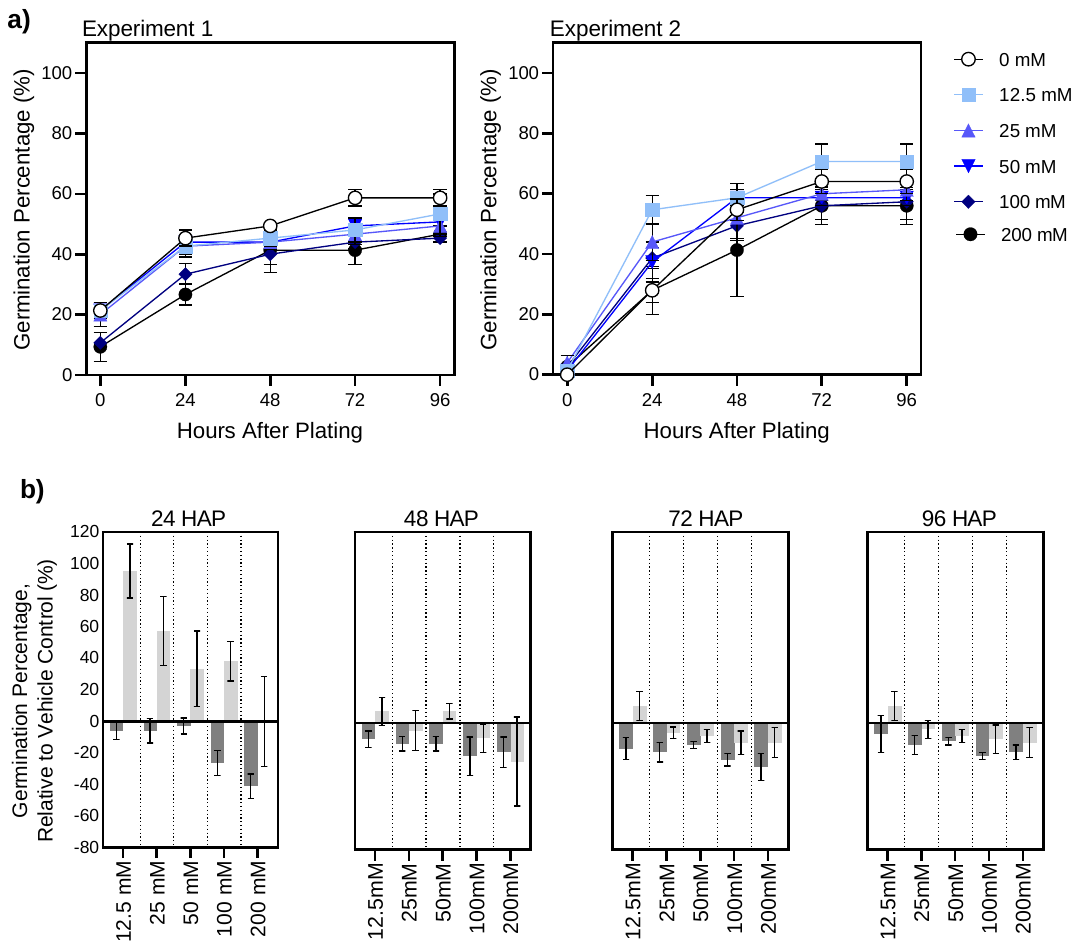

**Supplementary Figure 17**: Manganese (MnSO_4_) priming optimisation experiments in hemp (*Cannabis sativa*).

a) Germination percentage and b) germination percentage relative to the control of hemp seeds germinated in a 25^o^C, 16/8 h light/dark controlled incubator for 96 h. The data shown is the mean of 3 replicates and standard error is shown as vertical bars. In (b), dark grey and light grey refer to independent experiment 1 and 2 respectively. In (b) HAP refers to “hours after plating

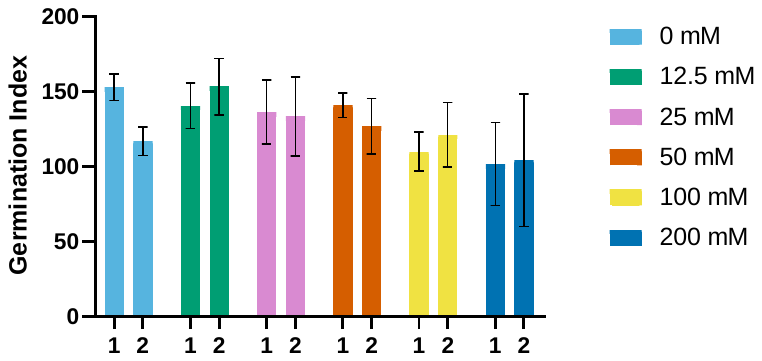

**Supplementary Figure 18**: Germination indices of hemp (*Cannabis sativa)* primed with varying concentrations of Manganese (MnSO_4_).

Hemp seeds were plated and grown in a 25^o^C, 16/8-h light/dark controlled incubator for 96 h. The data shown is the mean of 3 replicates and standard error is shown as vertical bars.

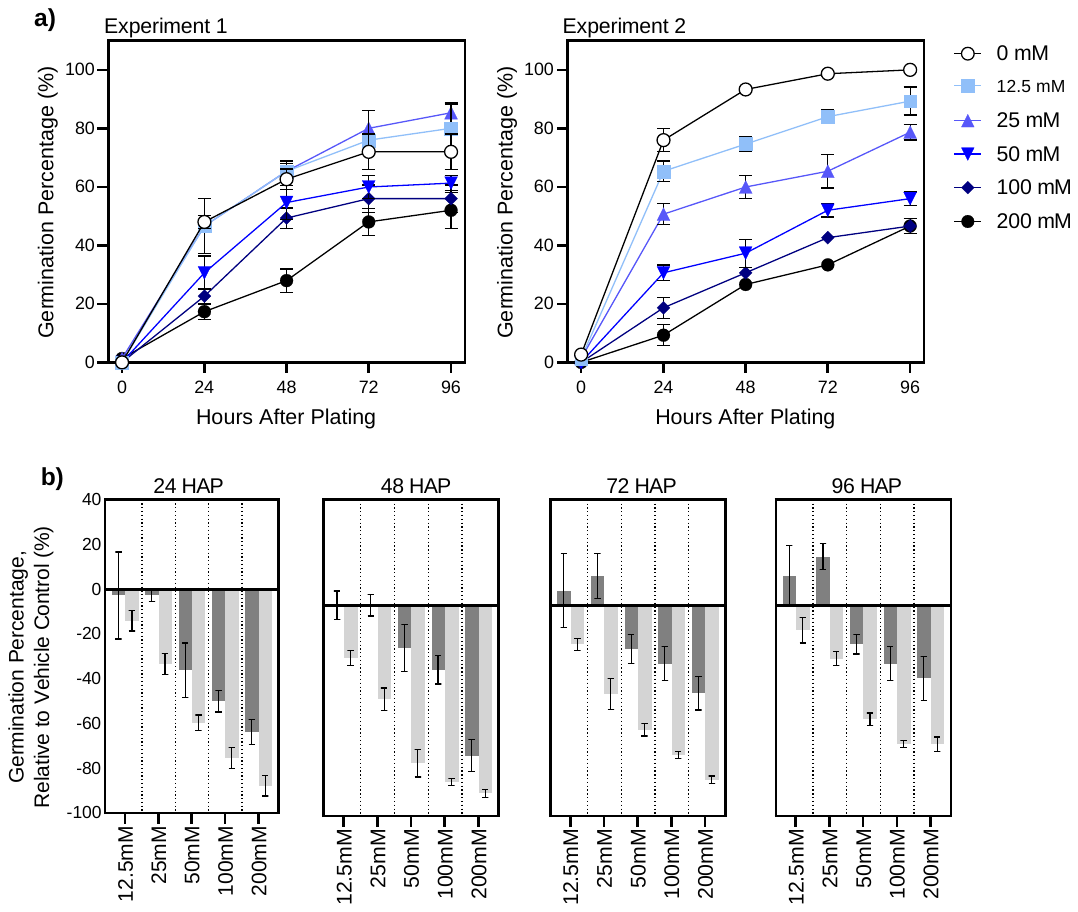

**Supplementary Figure 19**: Manganese (MnSO_4_) priming optimisation experiments in barley (*Hordeum vulgare*).

a) Germination percentage and b) germination percentage relative to the control of barley seeds germinated in a 25^o^C, 16/8-h light/dark controlled incubator for 96 h. The data shown is the mean of 3 replicates and standard error is shown as vertical bars. In (b), dark grey and light grey refer to independent experiment 1 and 2 respectively. In (b) HAP refers to “hours after plating”.

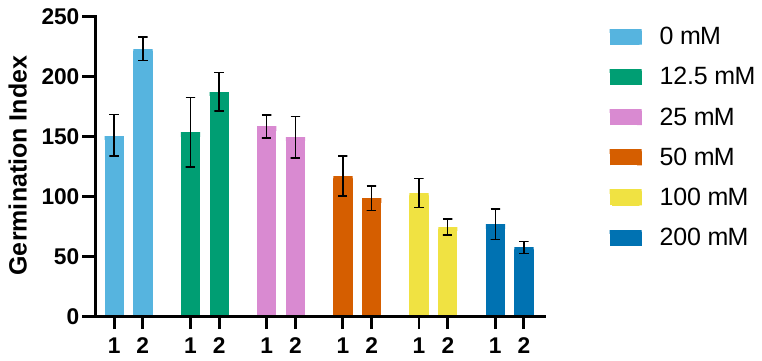

**Supplementary Figure 20**: Germination indices of barley (*Hordeum vulgare)* primed with varying concentrations of Manganese (MnSO_4_).

Hemp seeds were plated and grown in a 25^o^C, 16/8-h light/dark controlled incubator for 96 h. The data shown is the mean of 3 replicates and standard error is shown as vertical bars. “1” and “2” represent independent experiments done under the same conditions.

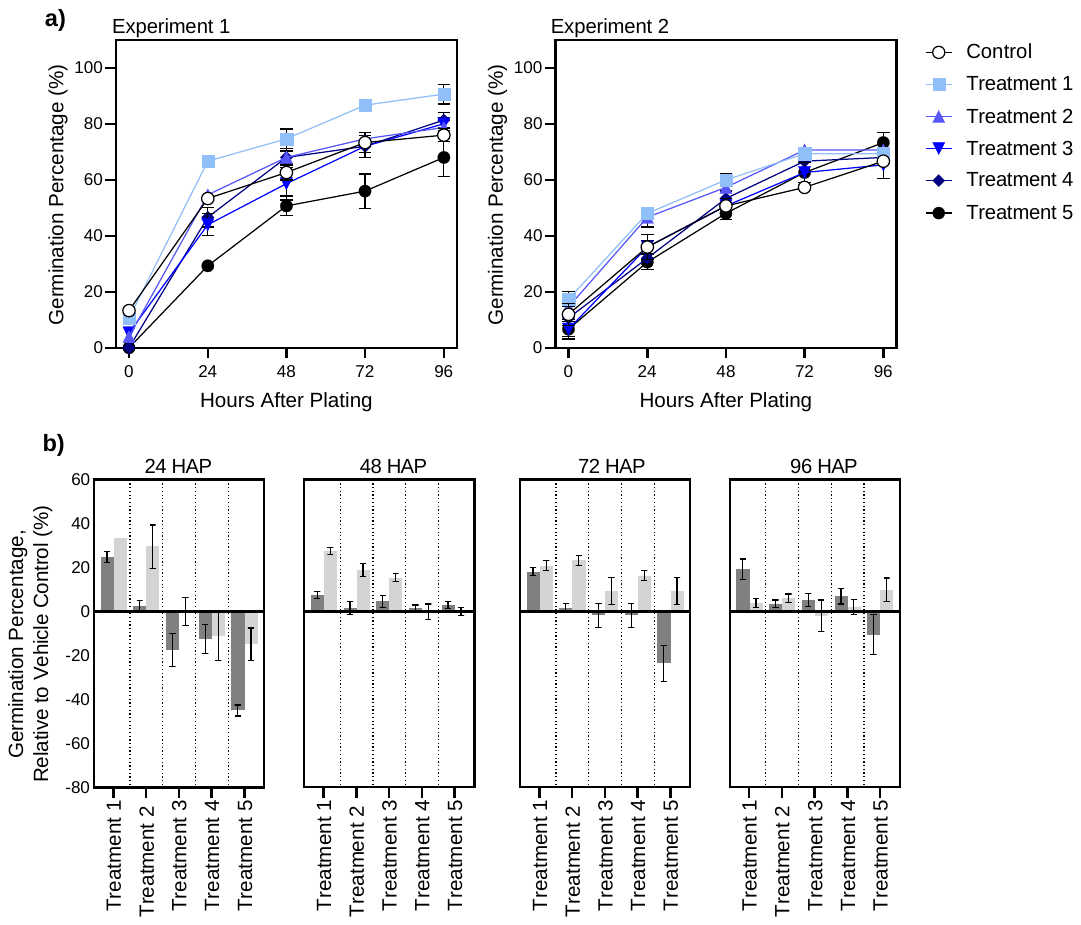

**Supplementary Figure 21**: Nutrient combination (ZnSO_4_, Na_2_SeO_3_, and MnSO_4_) priming optimisation experiments in hemp (*Cannabis sativa*).

a) Germination percentage and b) germination percentage relative to the control of hemp seeds germinated in a 25^o^C, 16/8-h light/dark controlled incubator for 96 h. Treatment 5: 100 mM ZnSO_4_, 50 µM Na_2_Se_3_O, 50 mM MnSO_4_, Treatment 4: 50 mM ZnSO_4_, 25 µM Na_2_Se_3_O, 25 mM MnSO_4_, Treatment 3: 25 mM ZnSO_4_, 12.5 µM Na_2_Se_3_O, 12.5 mM MnSO_4_, Treatment 2: 12.5 mM ZnSO_4_, 6.25 µM Na_2_Se_3_O, 6.25 mM MnSO_4_, Treatment 1: 6.25 mM ZnSO_4_, 3.125 µM Na_2_Se_3_O, 3.125 mM MnSO_4_. The data shown is the mean of 3 replicates and standard error is shown as vertical bars. In (b), dark grey and light grey refer to independent experiment 1 and 2 respectively. In (b) HAP refers to “hours after plating”.

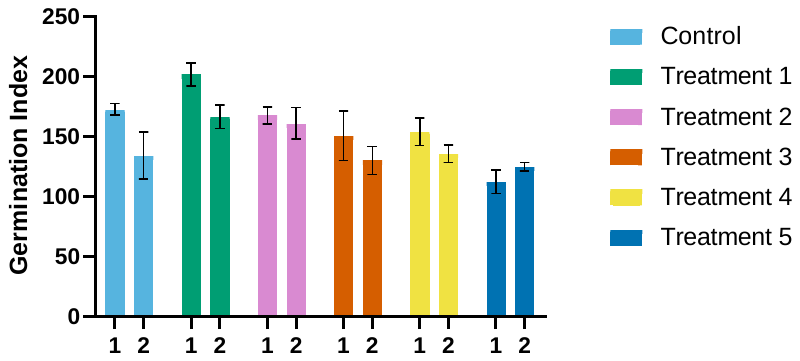

**Supplementary Figure 22**: Germination indices of Hemp (*Cannabis sativa)* primed with varying concentrations of micronutrients (ZnSO_4_, Na_2_SeO_3_, and MnSO_4_).

Hemp seeds were plated and grown in a 25^o^C, 16/8-h light/dark controlled incubator for 96 h. Treatment 5: 100 mM ZnSO_4_, 50 µM Na_2_Se_3_O, 50 mM MnSO_4_, Treatment 4: 50 mM ZnSO_4_, 25 µM Na_2_Se_3_O, 25 mM MnSO_4_, Treatment 3: 25 mM ZnSO_4_, 12.5 µM Na_2_Se_3_O, 12.5 mM MnSO_4_, Treatment 2: 12.5 mM ZnSO_4_, 6.25 µM Na_2_Se_3_O, 6.25 mM MnSO_4_, Treatment 1: 6.25 mM ZnSO_4_, 3.125 µM Na_2_Se_3_O, 3.125 mM MnSO_4_. The data shown is the mean of 3 replicates and standard error is shown as vertical bars. “1” and “2” represent independent experiments done under the same conditions.

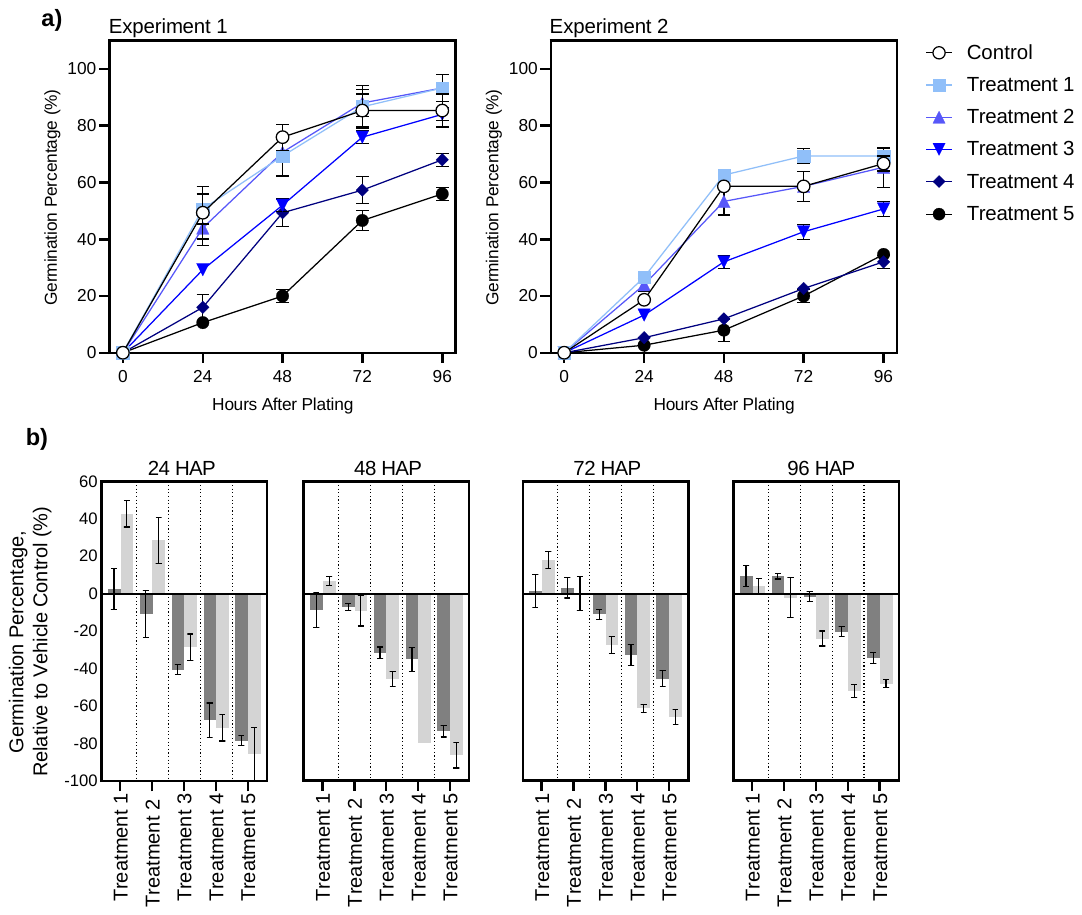

**Supplementary Figure 23**: Nutrient combination (ZnSO_4_, Na_2_SeO_3_, and MnSO_4_) priming optimisation experiments in barley (*Hordeum Vulgare*).

a) Germination percentage and b) germination percentage relative to the control of hemp seeds germinated in a 25^o^C, 16/8-h light/dark controlled incubator for 96 h. Treatment 5: 100mM ZnSO_4_, 50µM Na_2_Se_3_O, 50mM MnSO_4_, Treatment 4: 50 mM ZnSO_4_, 25 µM Na_2_Se_3_O, 25 mM MnSO_4_, Treatment 3: 25 mM ZnSO_4_, 12.5 µM Na_2_Se_3_O, 12.5 mM MnSO_4_, Treatment 2: 12.5 mM ZnSO_4_, 6.25 µM Na_2_Se_3_O, 6.25 mM MnSO_4_, Treatment 1: 6.25 mM ZnSO_4_, 3.125 µM Na_2_Se_3_O, 3.125 mM MnSO_4_. The data shown is the mean of 3 replicates and standard error is shown as vertical bars. In (b), dark grey and light grey refer to independent experiment 1 and 2 respectively. In (b) HAP refers to “hours after plating”.

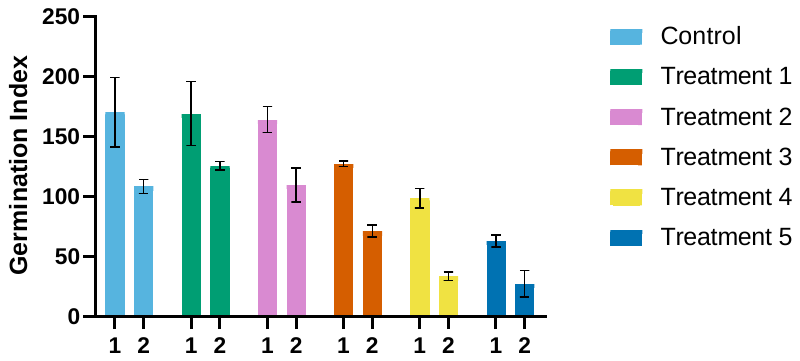

**Supplementary Figure 24**: Germination indices of barley (*Hordeum vulgare*) primed with varying concentrations of micronutrients (ZnSO_4_, Na_2_SeO_3_, and MnSO_4_).

Barley seeds were plated and grown in a 25^o^C, 16/8-h light/dark controlled incubator for 96 h. Treatment 5: 100 mM ZnSO_4_, 50 µM Na_2_Se_3_O, 50 mM MnSO_4_, Treatment 4: 50 mM ZnSO_4_, 25 µM Na_2_Se_3_O, 25 mM MnSO_4_, Treatment 3: 25 mM ZnSO_4_, 12.5 µM Na_2_Se_3_O, 12.5 mM MnSO_4_, Treatment 2: 12.5 mM ZnSO_4_, 6.25 µM Na_2_Se_3_O, 6.25 mM MnSO_4_, Treatment 1: 6.25 mM ZnSO_4_, 3.125 µM Na_2_Se_3_O, 3.125 mM MnSO_4_. The data shown is the mean of 3 replicates and standard error is shown as vertical bars. “1” and “2” represent independent experiments done under the same conditions.

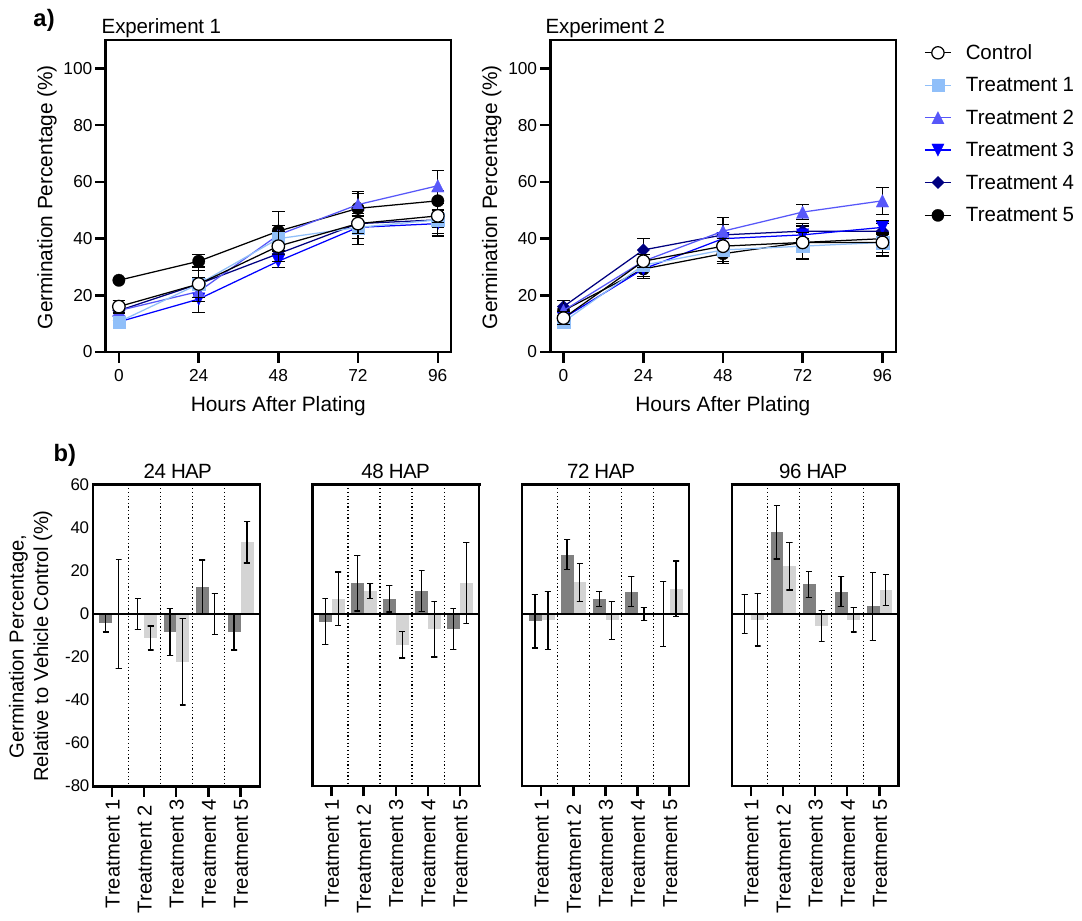

**Supplementary Figure 25**: Nutrient combination (ZnSO_4_, Na_2_SeO_3_, and MnSO_4_) priming optimisation experiments in hemp (*Cannabis sativa*).

a) Germination percentage and b) germination percentage relative to the control of hemp seeds germinated in a 25^o^C, 16/8-h light/dark controlled incubator for 96 h. Treatment 5: 25 mM ZnSO_4_, 12.5 µM Na_2_Se_3_O, 12.5 mM MnSO_4_. Treatment 4: 12.5 mM ZnSO_4_, 6.25 µM Na_2_Se_3_O, 6.25 mM MnSO_4_. Treatment 3: 6.25 mM ZnSO_4_, 3.125 µM Na_2_Se_3_O, 3.125 mM MnSO_4_. Treatment 2: 3.125 mM ZnSO_4_, 1.5 µM Na_2_Se_3_O, 1.5 mM MnSO_4_. Treatment 1: 1.5 mM ZnSO_4_, 0.75 µM Na_2_Se_3_O, 0.75 mM MnSO_4_. The data shown is the mean of 3 replicates and standard error is shown as vertical bars. In (b), dark grey and light grey refer to independent experiment 1 and 2 respectively. In (b) HAP refers to “hours after plating”.

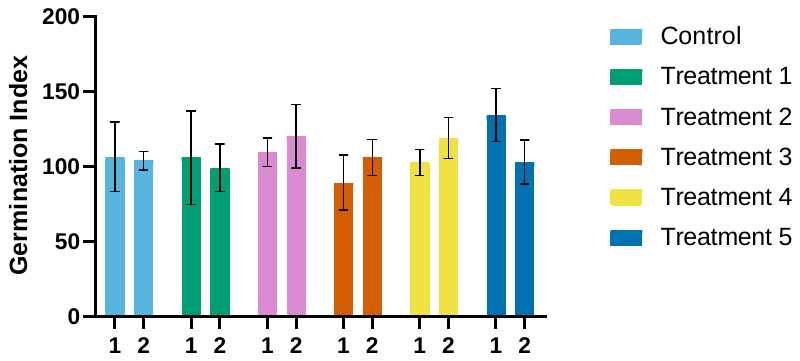

**Supplementary Figure 26**: Germination indices of hemp (*Cannabis sativa*) primed with varying concentrations of micronutrients (ZnSO_4_, Na_2_SeO_3_, and MnSO_4_).

Hemp seeds were plated and grown in a 25^o^C, 16/8-h light/dark controlled incubator for 96 h. Treatment 5: 25 mM ZnSO_4_, 12.5 µM Na_2_Se_3_O, 15.5mM MnSO_4_. Treatment 4: 12.5 mM ZnSO_4_, 6.25 µM Na_2_Se_3_O, 6.25 mM MnSO_4_. Treatment 3: 6.25 mM ZnSO_4_, 3.125 µM Na_2_Se_3_O, 3.125 mM MnSO_4_. Treatment 2: 3.125 mM ZnSO_4_, 1.5 µM Na_2_Se_3_O, 1.5 mM MnSO_4_. Treatment 1: 1.5 mM ZnSO_4_, 0.75 µM Na_2_Se_3_O, 0.75 mM MnSO_4_. The data shown is the mean of 3 replicates and standard error is shown as vertical bars. “1” and “2” represent independent experiments done under the same conditions.

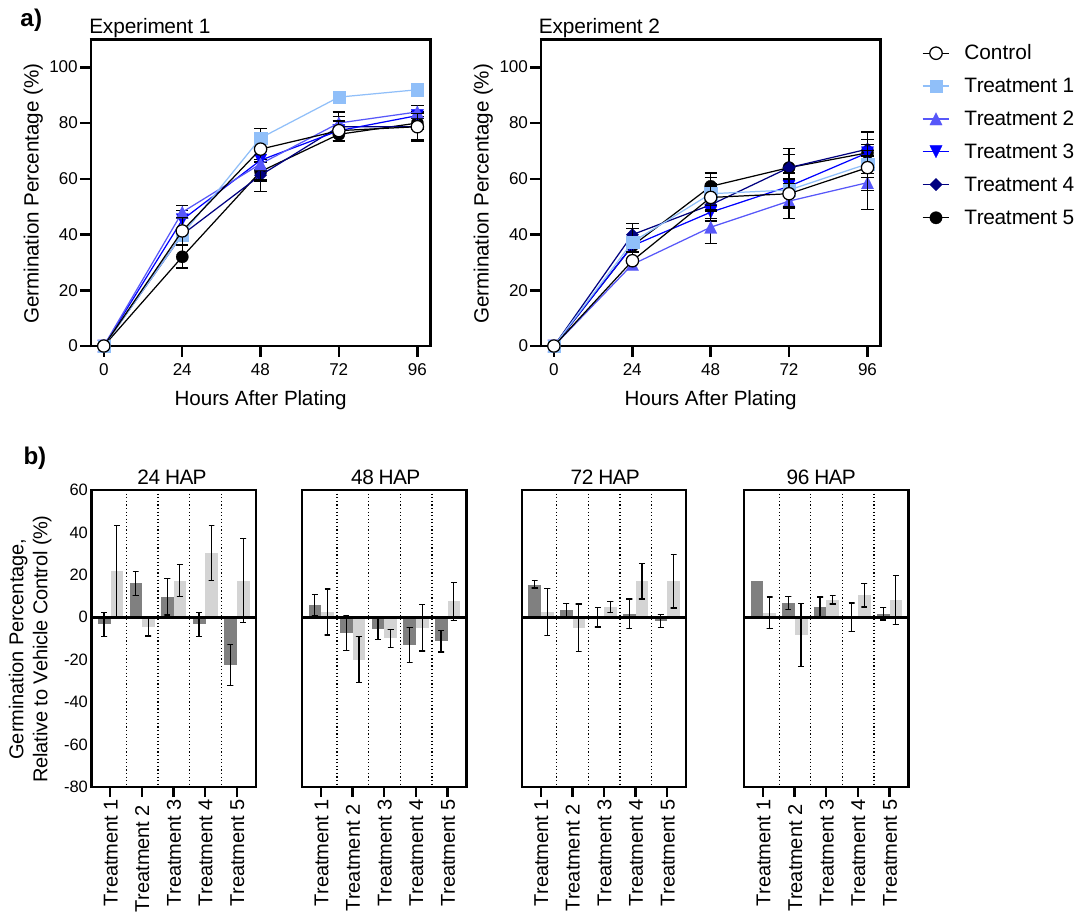

**Supplementary Figure 27**: Nutrient combination (ZnSO_4_, Na_2_SeO_3_, and MnSO_4_) priming optimisation experiments in barley (*Hordeum vulgare*).

a) Germination percentage and b) germination percentage relative to the control of barley seeds germinated in a 25^o^C, 16/8-h light/dark controlled incubator for 96 h. Treatment 5: 25 mM ZnSO_4_, 12.5 µM Na_2_Se_3_O, 12.5 mM MnSO_4_. Treatment 4: 12.5 mM ZnSO_4_, 6.25 µM Na_2_Se_3_O, 6.25 mM MnSO_4_. Treatment 3: 6.25 mM ZnSO_4_, 3.125 µM Na_2_Se_3_O, 3.125 mM MnSO_4_. Treatment 2: 3.125 mM ZnSO_4_, 1.5 µM Na_2_Se_3_O, 1.5 mM MnSO_4_. Treatment 1: 1.5 mM ZnSO_4_, 0.75 µM Na_2_Se_3_O, 0.75 mM MnSO_4_. The data shown is the mean of 3 replicates and standard error is shown as vertical bars. In (B), dark grey and light grey refer to independent experiment 1 and 2 respectively. In (B) HAP refers to “hours after plating”.

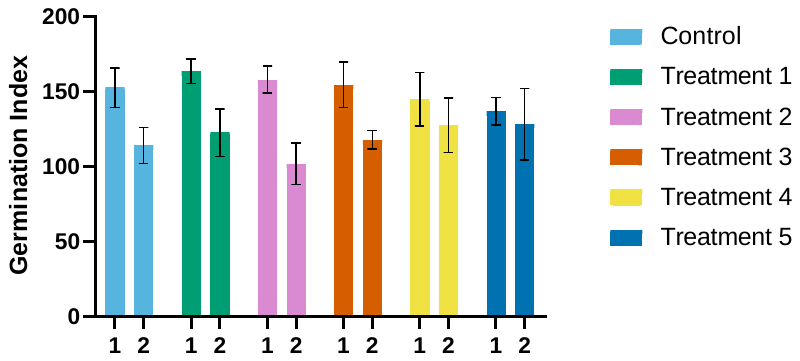

**Supplementary Figure 28**: Germination indices of barley (*Hordeum vulgare)* primed with varying concentrations of micronutrients (ZnSO_4_, Na_2_SeO_3_, and MnSO_4_).

Barley seeds were plated and grown in a 25^o^C, 16/8-h light/dark controlled incubator for 96 h. Treatment 5: 25 mM ZnSO_4_, 12.5 µM Na_2_Se_3_O, 12.5 mM MnSO_4_. Treatment 4: 12.5 mM ZnSO_4_, 6.25 µM Na_2_Se_3_O, 6.25 mM MnSO_4_. Treatment 3: 6.25 mM ZnSO_4_, 3.125 µM Na_2_Se_3_O, 3.125 mM MnSO_4_. Treatment 2: 3.125 mM ZnSO_4_, 1.5 µM Na_2_Se_3_O, 1.5 mM MnSO_4_. Treatment 1: 1.5 mM ZnSO_4_, 0.75 µM Na_2_Se_3_O, 0.75 mM MnSO_4_. The data shown is the mean of 3 replicates and standard error is shown as vertical bars. “1” and “2” represent independent experiments done under the same conditions.

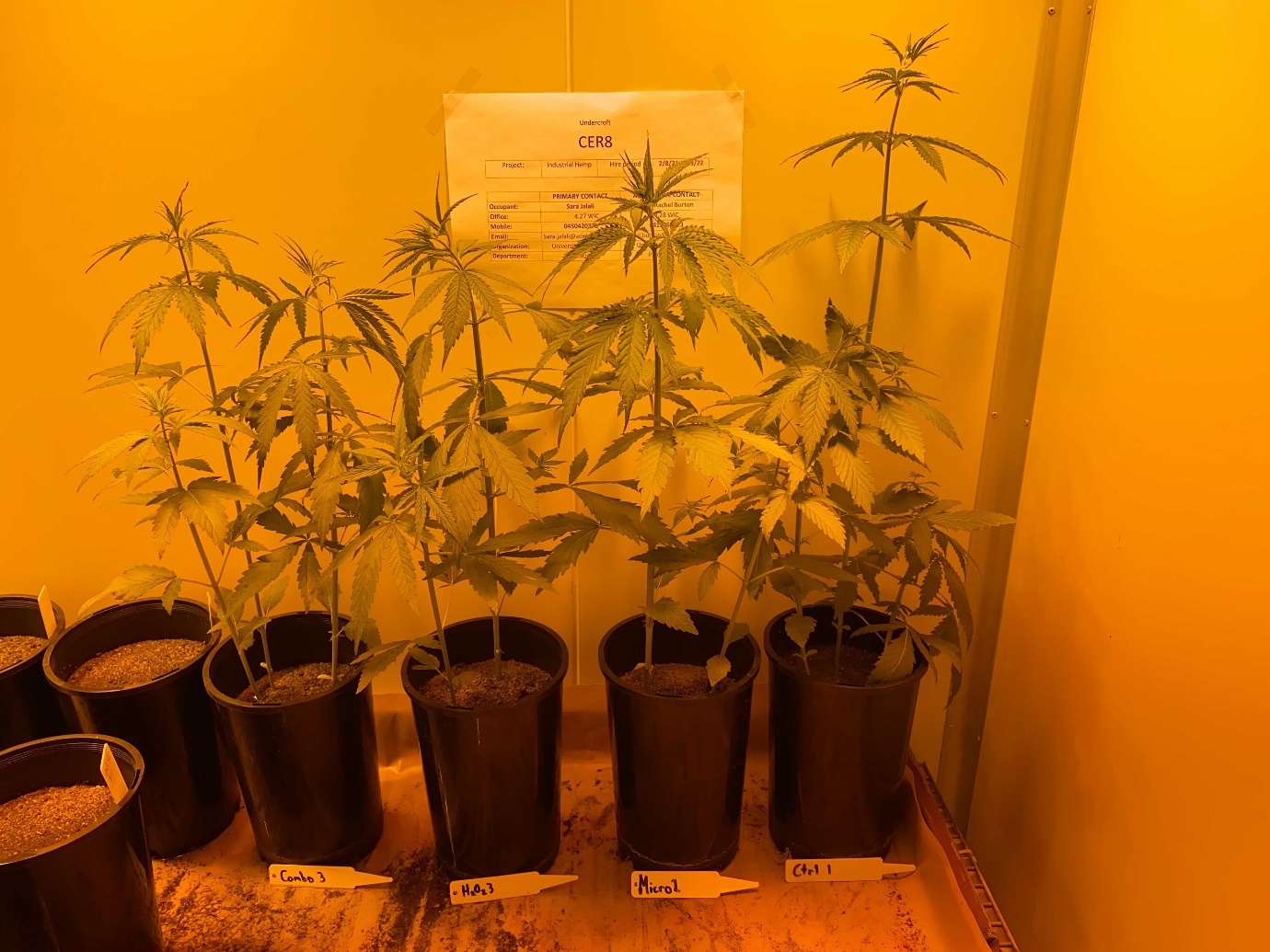

Control

Micronutrient

H_2_O_2_

Combination

**Supplementary Figure 29**: The final time point (three weeks) for the potted growth of primed hemp seeds.

Hemp seeds were primed under different treatments (as) labelled potted in soil grown in 16/8 hr light/dark photoperiod, 18/26 °C night/day controlled environment room for 3 weeks before harvest. Nutrient Treatment: 12.5 mM ZnSO_4_, 6.25 µM Na_2_Se_3_O, 6.25 mM MnSO_4_. H_2_O_2_ Treatment: 125 mM H_2_O_2_. Combination Treatment: 125 mM H_2_O_2_, 12.5 mM ZnSO_4_, 6.25 µM Na_2_Se_3_O, 6.25 mM MnSO_4_

For scale, the pots are 15 cm in diameter.

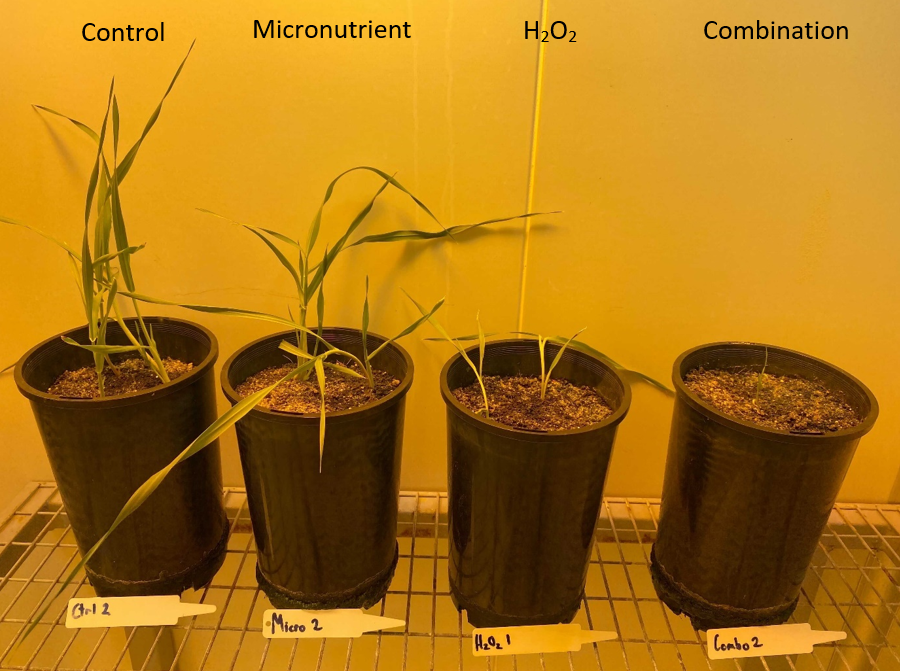

**Supplementary Figure 30**: The final time point (three weeks) for the potted growth of primed barley seeds.

Barley seeds were primed under different treatments (as) labelled potted in soil grown in 16/8 hr light/dark photoperiod, 18/23 °C night/day-controlled environment room for 3 weeks before harvest. Nutrient Treatment: 12.5 mM ZnSO_4_, 6.25 µM Na_2_Se_3_O, 6.25 mM MnSO_4_. H_2_O_2_ Treatment: 125 mM H_2_O_2_. Combination Treatment: 125 mM H_2_O_2_, 12.5 mM ZnSO_4_, 6.25 µM Na_2_Se_3_O, 6.25 mM MnSO_4_.

For scale, the pots are 15 cm in diameter.
